# Temporal transcriptional response of *Candida glabrata* during macrophage infection reveals a multifaceted transcriptional regulator CgXbp1 important for macrophage response and drug resistance

**DOI:** 10.1101/2021.09.28.462173

**Authors:** Maruti Nandan Rai, Chirag Parsania, Rikky Rai, Niranjan Shirgaonkar, Kaeling Tan, Koon Ho Wong

**Author notes:** For correspondence, please contact. Rikky Rai –.

## Abstract

*Candida glabrata* can thrive inside macrophages and tolerate high levels of azole antifungals. These innate abilities render infections by this human pathogen a clinical challenge. How *C. glabrata* reacts inside macrophages and what is the molecular basis of its drug tolerance are not well understood. Here, we mapped genome-wide RNA polymerase II (RNAPII) occupancy in *C. glabrata* to delineate its transcriptional responses during macrophage infection in high temporal resolution. RNAPII profiles revealed dynamic *C. glabrata* responses to macrophage with genes of specialized pathways activated chronologically at different times of infection. We identified an uncharacterized transcription factor (CgXbp1) important for the chronological macrophage response, survival in macrophages, and virulence. Genome-wide mapping of CgXbp1 direct targets further revealed its multi-faceted functions, regulating not only virulence-related genes but also genes associated with drug resistance. Finally, we showed that CgXbp1 indeed also affects azole resistance. Overall, this work presents a powerful approach for examining host-pathogen interaction and uncovers a novel transcription factor important for *C. glabrata*’s survival in macrophages and drug tolerance.

## Introduction

Phagocytes such as macrophages constitute the first line of host immune defence against invading pathogens (Brown, 2011; Erwig and Gow, 2016). The ability to escape or survive phagocytic attacks is fundamental to the virulence of pathogens (Seider *et al*, 2010; Erwig & Gow, 2016). *Candida* species are prominent opportunistic fungal pathogens with an associated mortality rate of ∼29-60% among immunocompromised population (Bongomin *et al*., 2017; Lamoth *et al*., 2018). *Candida albicans* is responsible for most Candidiasis infections, although recent studies indicate an epidemiological shift in Candidiasis with an upsurge in infections caused by *Candida glabrata* (Benedict *et al*, 2017; Lamoth *et al*, 2018). Relative to other fungal species including *C. albicans*, *C. glabrata* is more resistant to antifungal drugs like fluconazole and can survive and proliferate inside immune cells (Seider *et al*., 2011; Rai *et al*., 2012). Thus far, details about how *C. glabrata* survives, adapts and proliferates in phagocytes and the basis for its intrinsically high azole resistance are still not clearly understood.

Genome-wide transcriptomic studies have been performed to map the response of *Candida species* during macrophage infection (Lorenz & Fink, 2001; Rubin-Bejerano *et al*, 2003; Lorenz *et al*, 2004; Kaur *et al*, 2007; Rai *et al*, 2012), but the insights gained into the infection process so far lack temporal resolution, centring mostly on the late stages of the pathogen-host interactions. We reason that the immediate and early pathogen response is pivotal for survival and adaptation in the host, while responses during later stages reflect strategies for growth and proliferation. Therefore, delineating the whole episode of pathogen response, instead of just a snapshot, during infection is fundamental to understanding pathogenesis. However, conventional transcriptomic analysis involving mRNAs are not suitable for dissecting dynamic temporal transcriptional changes, as measurements of mRNA levels are convoluted by transcript stabilities (Tan and Wong, 2019).

Here, we applied the powerful Chromatin Immuno-precipitation followed by Next Generation Sequencing (ChIP-seq) method against elongating RNA Polymerase II (RNAPII) to map *C. glabrata* transcriptional responses during macrophage infection. We show that *C. glabrata* responds to macrophage infection by mounting chronological transcriptional responses. Based on the expression pattern, we identified many candidate transcriptional regulators including a novel transcription factor, CgXbp1, for the macrophage response. Deletion of CgXbp1 led to accelerated transcriptional activation of genes associated with multiple biological processes during interaction with macrophages. We further demonstrate that CgXbp1 is a multifaceted transcription factor directly binding to ∼10% of *C. glabrata* genes including many involves in the pathogenesis process and drug resistance. *CgXBP1* deletion resulted in attenuated survival in host macrophages, diminished virulence in the *Galleria mellonella* model of Candidiasis, and elevated resistance to antifungal drug fluconazole. Overall, our work uncovers an important novel transcription factor for *C. glabrata*’s survival in macrophages and antifungal drug resistance.

## Results

### Mapping high temporal resolution transcriptional responses of *C. glabrata* during macrophage infection

To understand how *C. glabrata* survives macrophage phagocytosis, we applied ChIP-seq against elongating RNAPII in a time-course experiment after 0.5, 2, 4, 6, and 8 h of THP-1 macrophage infection to map genome-wide transcription responses of *C. glabrata* during different stages of THP-1 macrophage infection (Figure 1A). As expected, genes known to be induced by macrophage phagocytosis (e.g. tricarboxylic acid [TCA] cycle, glyoxylate bypass, and iron homeostasis genes (Kaur *et al*, 2007; Rai *et al*, 2012)) had significant RNAPII occupancies at their gene bodies specifically but not at inter-genic regions (Figure 1B, Figure 1-figure supplement 1A). In addition, the ChIP-seq data also revealed temporal gene expression information. For example, the ATP synthesis gene *CgCYC1* was dramatically up-regulated immediately (0.5 h) upon macrophage internalisation, while the TCA cycle gene *CgCIT2* and glyoxylate bypass gene *CgICL1* were induced slightly later at 2 h and their transcription levels decreased subsequently (4-6 h) (Figure 1B). In contrast, an opposite transcription pattern (e.g. gradual increasing and peaking at later stages) was observed for *CgFTR1*, *CgTRR1,* and *CgMT-I*, which are involved in iron uptake, biofilm formation, and sequestration of metal ions, respectively (Figure 1B). Therefore, the RNAPII ChIP-seq approach can effectively capture a high temporal resolution gene expression landscape in *C. glabrata* during macrophage infection.

**Figure 1:**
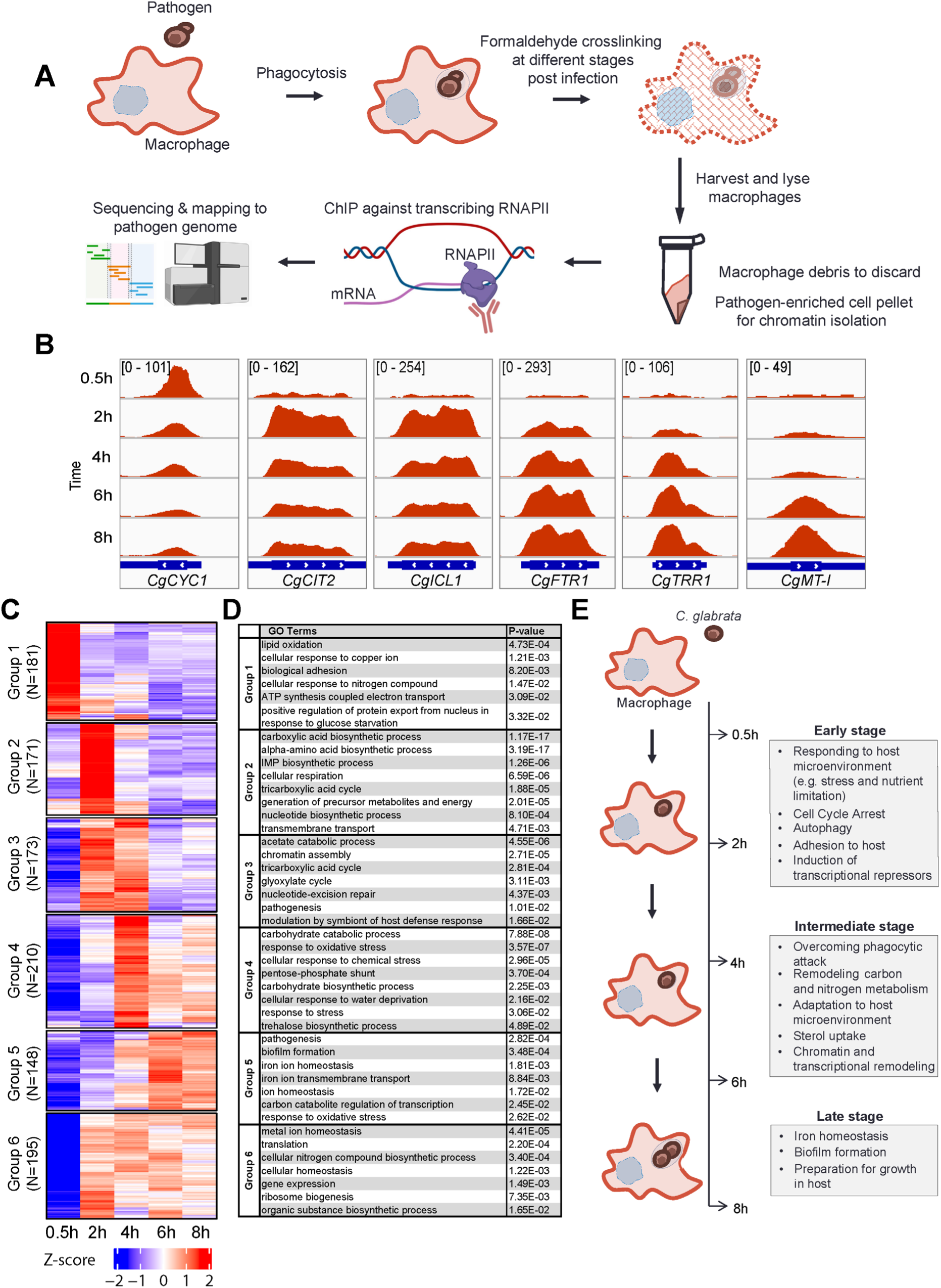
*C. glabrata* mounts a dynamic chronological transcriptional response upon macrophage infection. **(A)** A schematic diagram showing the overall methodology used in this study. **(B)** Genome browser views of RNAPII ChIP-seq signals on *CgCYC1*, *CgCIT2*, *CgICL1*, *CgFTR1*, *CgTRR1* and *CgMT-I* genes at indicated time points. Numbers in the square brackets indicate the y-axis scale range of normalized RNAPII ChIP-seq signal used for the indicated genes across different datasets. **(C)** A heatmap showing temporal expression patterns of transcribed genes in *C. glabrata* during 0.5 to 8 h macrophage infection in a time-course experiment. The colour scale represents the Z-score of the normalized RNAPII ChIP-seq signal. **(D)** A table showing significantly enriched GO biological processes (P value ≤ 0.05) for the six groups of temporally transcribed genes. **(E)** A schematic diagram showing *C. glabrata* transcriptional responses (broadly classified into early, intermediate and late stages) during macrophage infection.

### *C. glabrata* mounts dynamic, temporal and chronological transcription responses during macrophage infection

Systematic analysis of actively transcribed genes revealed that approximately 30% of *C. glabrata* genes (n = 1,589, Supplementary File 1) were constitutively transcribed (n = 511, Figure 1-figure supplement 1B) or temporally induced (n = 1,078, Figure 1-figure supplement 1C) during macrophage infection. Temporally induced genes were further classified into six groups according to their transcription pattern using *k*-means clustering. The overall transcriptional response was highly diverse with each group exhibiting a unique temporal transcriptional pattern (Figure 1C). Interestingly, while some genes were induced immediately (0.5 h, Group 1, n = 181) upon internalisation by macrophages, transcriptional induction of over 80% of genes (Group 2-6, n = 897) did not happen until later (2-8 h). Besides, their expression patterns were highly variable, illustrating the complex and dynamic nature of *C. glabrata* transcriptional response during macrophage infection.

Gene Ontology (GO) analysis revealed chronological activation of different biological processes during the infection process (Figure 1D, Supplementary File 2). In the immediate response (0.5 h), genes (Group 1, n = 181) were significantly enriched in processes such as adhesion, responses to copper ion and nitrogen compound, positive regulation of nuclear export in response to glucose starvation, lipid oxidation, and ATP synthesis (Figure 1D). This indicated that *C. glabrata* experiences nutrient and energy deprivation immediately upon entry to macrophages (Figure 1E). Alternatively, the induction of ATP biosynthesis genes may reflect a strong demand for energy by *C. glabrata* to deal with the host’s attacks and/or to adapt to the host microenvironment. Subsequently (2 h post phagocytosis), *C. glabrata* underwent a major metabolic remodelling presumably to prepare for growth and generate energy, as reflected by the next wave of transcriptional induction for genes (Group 2, n = 171) involved in the TCA cycle, biosynthesis of inosine 5’ monophosphate (IMP), carboxylic acid, amino acid, nucleotide, and precursor for metabolite and energy (Figures 1C&D). In addition, cell cycle arrest and DNA damage checkpoint genes (*CgMEC3, CgGLC7, CAGL0G07271g, and CAGL0A04213g*) were also strongly induced at this early stage, and *C. glabrata* cells were indeed arrested at the G1-S phase cell cycle after macrophage engulfment (Figure 1-figure supplement 2). It is noteworthy that many genes and pathways previously shown to be critical for *C. glabrata* virulence (Kaur *et al*., 2005; Rai *et al*., 2012; Kasper *et al*., 2015) such as adherence, response to DNA damage, oxidative stress, autophagy, TCA cycle, amino acid biosynthesis, and iron homeostasis (Figure 1-figure supplement 3A-G), were markedly induced at the early stages (0.5 and 2 h). Therefore, virulence-centric biological processes were among the most immediate *C. glabrata* responses upon macrophage phagocytosis (Figure 1E), implying the importance of the early transcriptional response towards its adaptation and survival in macrophages.

During the next stage of infection (2-4 h), *C. glabrata* continued to actively transcribe genes associated with carbon metabolism, DNA repair and pathogenesis (Group 3, Figures 1C&D), suggesting that the invading pathogen was still trying to achieve metabolic homeostasis and to counter macrophage internal milieu. Genes required for chromatin assembly and modification were also significantly induced at this stage (Figure 1-figure supplement 4A&B), supporting an earlier report about the involvement of chromatin remodelling during the infection process (Rai *et al*., 2012). Towards the later phase of this stage (4 h), genes for responses to different stresses (e.g. oxidative, chemical stress, and osmolarity) and resistance thermo-tolerance and oxidative stress (e.g. trehalose biosynthesis and pentose phosphate pathway, respectively) become maximally induced (Group 4, Figures 1C&D). The induction of these stress response pathway genes towards the end of metabolic remodelling is somewhat unexpected, as it suggests that phagocytic attacks (e.g. ROS) against *C. glabrata* might not have occurred until the later phase. However, as shown above, the observations that DNA repair and damage response genes were already upregulated at 2 h indicate that cells had already experienced the attacks. These findings collectively suggest that *C. glabrata* elicits a coordinated stage-wise response during infection; first adapting to macrophage nutrient microenvironment before overcoming phagocytic attacks (Figure 1E). Interestingly, a family of sterol uptake genes (known as *TIR* [Tip1-related]) displayed concerted transcription activation at the end of this stage (4 h) (Figure 1-figure supplement 5). In *Saccharomyces cerevisiae*, *TIR* genes are activated and required for growth under anaerobic condition (Abramova *et al*., 2001). Given that sterols are an essential component of the cell membrane and that ergosterol biosynthesis is an oxygen-dependent process (Joffrion and Cushion, 2010), the up-regulation of the TIR genes indicates an experience of oxygen deprivation and a need for sterols (presumably for proliferation) by *C. glabrata*.

Towards the late stage (6-8 h), genes required for biofilm formation, iron homeostasis, both of which play critical roles in the pathogenesis process (Seider *et al*, 2014; Rodrigues *et al*, 2017), became maximal induced (Group 5, Figures 1C&D). As biofilm formation involves cell growth and proliferation, this observation potentially suggests that the cells are preparing for growth, and this is consistent with the concomitant induction of the iron homeostasis genes that are also necessary for proliferation. Altogether, the overall results revealed details into the dynamic stage-wise responses of *C. glabrata* during macrophage infection (Figure 1E).

### Identification of potential transcriptional regulators of early temporal response

We next attempted to identify the potential transcriptional regulators for the chronological transcriptional response. Remarkably, more than 25% of *C. glabrata* transcription factor (TF) genes (n = 53) were expressed during macrophage infection (Table 1), with 39 TF genes showing a temporal induction pattern (Figure 2A). Of note, eleven TFs (Aft1, Ap1, Ap5, Haa1, Hap4, Hap5, Msn4, Upc2, Yap3, Yap6 and Yap7) are known to either bind or control some of the macrophage infection-induced genes (Supplementary File 3) as reported by PathoYeastract (Monteiro *et al*, 2020), providing strong support to the identified TFs being responsible for the observed temporal transcriptional response.

**Figure 2:**
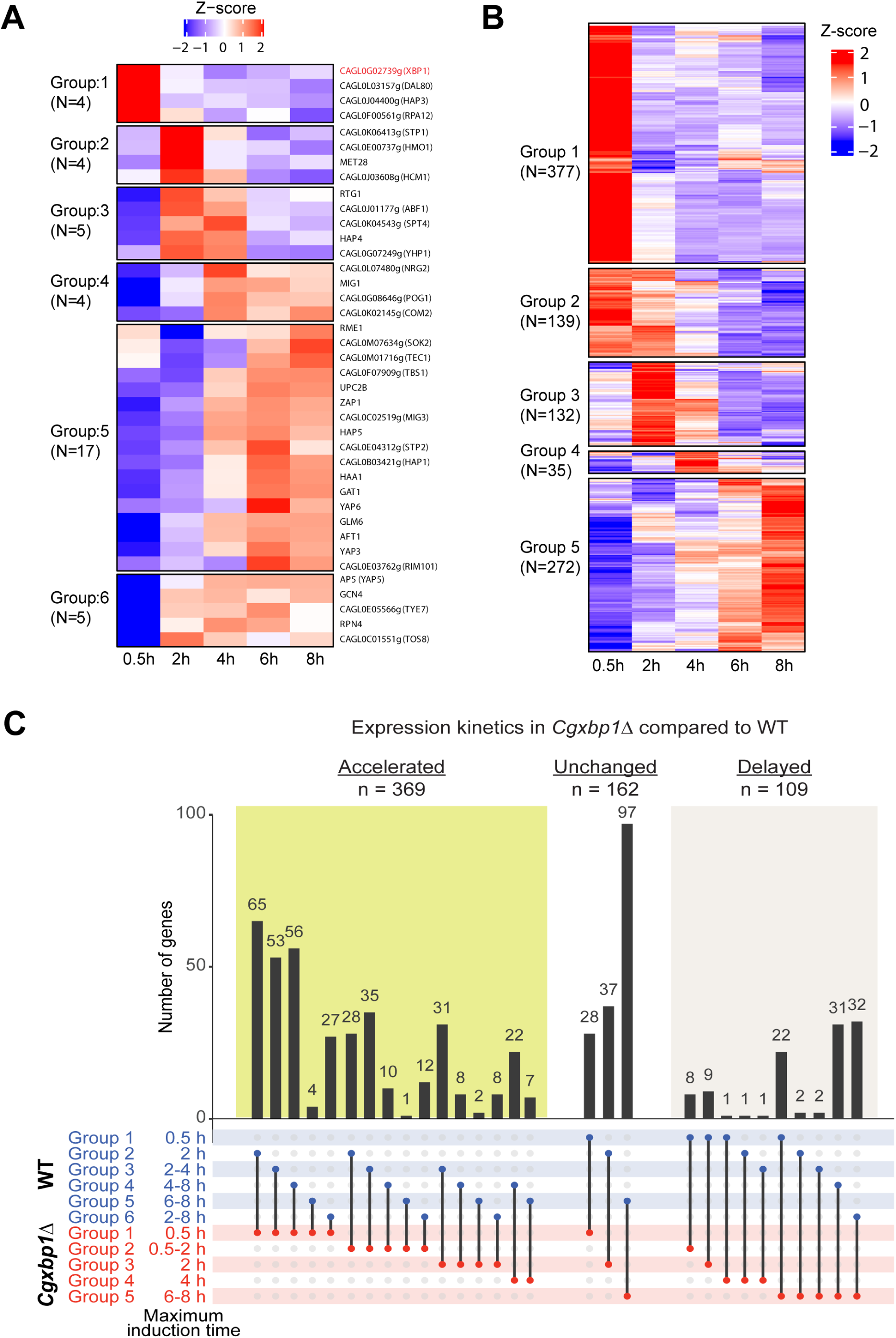
CgXbp1 is central in orchestrating the dynamic transcriptional response of *C. glabrata* during macrophage infection. **(A)** A heatmap showing temporal expression patterns of *C. glabrata* transcription factor genes transcribed during THP-1 macrophage infection. Colour scale represents the Z-score of the normalized RNAPII ChIP-seq signal. **(B)** A heatmap showing temporal expression patterns of transcribed genes in the *Cgxbp1*Δ mutant during 0.5 to 8 h THP-1 macrophage infection in a time-course experiment. **(C)** An UpSet plot showing the number of genes induced at the indicated time points in WT and the *Cgxbp1*Δ mutant during THP-1 macrophage infection.

**Table 1:**
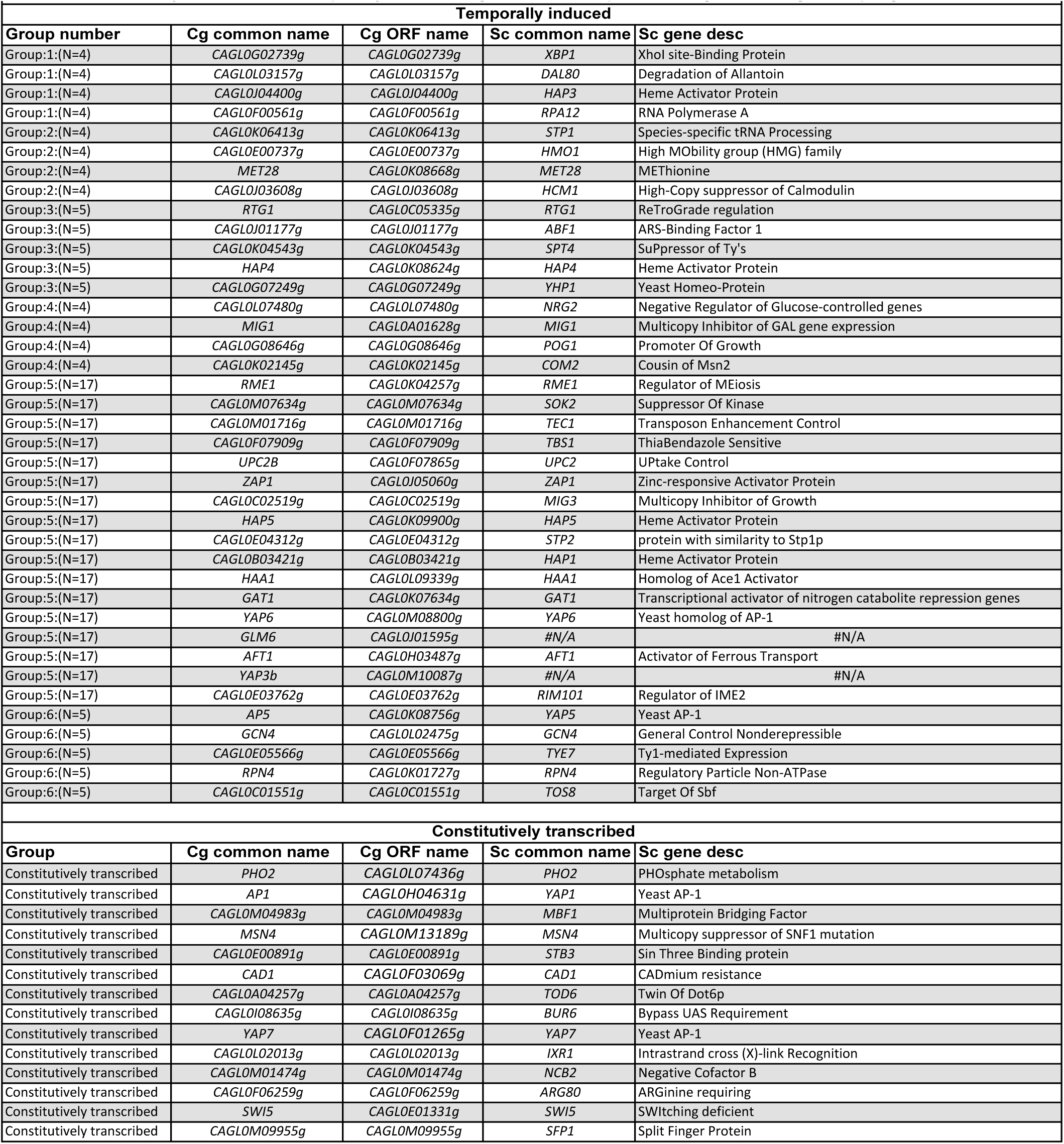
Constitutively transcribed or temporally induced *C. glabrata* transcription factor genes during macrophage infection.

|  |  |  |  |  |
| --- | --- | --- | --- | --- |
| Temporally induced |  |  |  |  |
| Group number | Cg common name | Cg ORF name | Sc common name | Sc gene desc |
| Group:1:(N=4) | CAGL0G02739g | CAGL0G02739g | XBP1 | XhoI site-Binding Protein |
| Group:1:(N=4) | CAGL0L03157g | CAGL0L03157g | DAL80 | Degradation of Allantoin |
| Group:1:(N=4) | CAGL0J04400g | CAGL0J04400g | HAP3 | Heme Activator Protein |
| Group:1:(N=4) | CAGL0F00561g | CAGL0F00561g | RPA12 | RNA Polymerase A |
| Group:2:(N=4) | CAGL0K06413g | CAGL0K06413g | STP1 | Species-specific tRNA Processing |
| Group:2:(N=4) | CAGL0E00737g | CAGL0E00737g | HMO1 | High MOBility group (HMG) family |
| Group:2:(N=4) | MET28 | CAGL0K08668g | MET28 | METHionine |
| Group:2:(N=4) | CAGL0J03608g | CAGL0J03608g | HCM1 | High-Copy suppressor of Calmodulin |
| Group:3:(N=5) | RTG1 | CAGL0C05335g | RTG1 | ReTroGrade regulation |
| Group:3:(N=5) | CAGL0J01177g | CAGL0J01177g | ABF1 | ARS-Binding Factor 1 |
| Group:3:(N=5) | CAGL0K04543g | CAGL0K04543g | SPT4 | SuPpressor of Ty's |
| Group:3:(N=5) | HAP4 | CAGL0K08624g | HAP4 | Heme Activator Protein |
| Group:3:(N=5) | CAGL0G07249g | CAGL0G07249g | YHP1 | Yeast Homeo-Protein |
| Group:4:(N=4) | CAGL0L07480g | CAGL0L07480g | NRG2 | Negative Regulator of Glucose-controlled genes |
| Group:4:(N=4) | MIG1 | CAGL0A01628g | MIG1 | Multicopy Inhibitor of GAL gene expression |
| Group:4:(N=4) | CAGL0G08646g | CAGL0G08646g | POG1 | Promoter Of Growth |
| Group:4:(N=4) | CAGL0K02145g | CAGL0K02145g | COM2 | Cousin of Msn2 |
| Group:5:(N=17) | RME1 | CAGL0K04257g | RME1 | Regulator of MEiosis |
| Group:5:(N=17) | CAGL0M07634g | CAGL0M07634g | SOK2 | Suppressor Of Kinase |
| Group:5:(N=17) | CAGL0M01716g | CAGL0M01716g | TEC1 | Transposon Enhancement Control |
| Group:5:(N=17) | CAGL0F07909g | CAGL0F07909g | TBS1 | ThiaBendazole Sensitive |
| Group:5:(N=17) | UPC2B | CAGL0F07865g | UPC2 | UPTake Control |
| Group:5:(N=17) | ZAP1 | CAGL0J05060g | ZAP1 | Zinc-responsive Activator Protein |
| Group:5:(N=17) | CAGL0C02519g | CAGL0C02519g | MIG3 | Multicopy Inhibitor of Growth |
| Group:5:(N=17) | HAP5 | CAGL0K09900g | HAP5 | Heme Activator Protein |
| Group:5:(N=17) | CAGL0E04312g | CAGL0E04312g | STP2 | protein with similarity to Stp1p |
| Group:5:(N=17) | CAGL0B03421g | CAGL0B03421g | HAP1 | Heme Activator Protein |
| Group:5:(N=17) | HAA1 | CAGL0L09339g | HAA1 | Homolog of Ace1 Activator |
| Group:5:(N=17) | GAT1 | CAGL0K07634g | GAT1 | Transcriptional activator of nitrogen catabolite repression genes |
| Group:5:(N=17) | YAP6 | CAGL0M08800g | YAP6 | Yeast homolog of AP-1 |
| Group:5:(N=17) | GLM6 | CAGL0J01595g | #N/A | #N/A |
| Group:5:(N=17) | AFT1 | CAGL0H03487g | AFT1 | Activator of Ferrous Transport |
| Group:5:(N=17) | YAP3b | CAGL0M10087g | #N/A | #N/A |
| Group:5:(N=17) | CAGL0E03762g | CAGL0E03762g | RIM101 | Regulator of IME2 |
| Group:6:(N=5) | AP5 | CAGL0K08756g | YAP5 | Yeast AP-1 |
| Group:6:(N=5) | GCN4 | CAGL0L02475g | GCN4 | General Control Nonderepressible |
| Group:6:(N=5) | CAGL0E05566g | CAGL0E05566g | TYE7 | Ty1-mediated Expression |
| Group:6:(N=5) | RPN4 | CAGL0K01727g | RPN4 | Regulatory Particle Non-ATPase |
| Group:6:(N=5) | CAGL0C01551g | CAGL0C01551g | TOS8 | Target Of Sbf |
| Constitutively transcribed |  |  |  |  |
| Group | Cg common name | Cg ORF name | Sc common name | Sc gene desc |
| Constitutively transcribed | PHO2 | CAGL0L07436g | PHO2 | PHOSphate metabolism |
| Constitutively transcribed | AP1 | CAGL0H04631g | YAP1 | Yeast AP-1 |
| Constitutively transcribed | CAGL0M04983g | CAGL0M04983g | MBF1 | Multiprotein Bridging Factor |
| Constitutively transcribed | MSN4 | CAGL0M13189g | MSN4 | Multicopy suppressor of SNF1 mutation |
| Constitutively transcribed | CAGL0E00891g | CAGL0E00891g | STB3 | Sin Three Binding protein |
| Constitutively transcribed | CAD1 | CAGL0F03069g | CAD1 | CADmium resistance |
| Constitutively transcribed | CAGL0A04257g | CAGL0A04257g | TOD6 | Twin Of Dot6p |
| Constitutively transcribed | CAGL0I08635g | CAGL0I08635g | BUR6 | Bypass UAS Requirement |
| Constitutively transcribed | YAP7 | CAGL0F01265g | YAP7 | Yeast AP-1 |
| Constitutively transcribed | CAGL0L02013g | CAGL0L02013g | IXR1 | Intrastrand cross (X)-link Recognition |
| Constitutively transcribed | CAGL0M01474g | CAGL0M01474g | NCB2 | Negative Cofactor B |
| Constitutively transcribed | CAGL0F06259g | CAGL0F06259g | ARG80 | ARGINine requiring |
| Constitutively transcribed | SWI5 | CAGL0E01331g | SWI5 | SWItching deficient |
| Constitutively transcribed | CAGL0M09955g | CAGL0M09955g | SFP1 | Split Finger Protein |

As the early response is likely to have influential effects on infection outcome, we focused on the four candidate TFs in Group 1 (induction at 0.5 h); the genes *CAGL0F00561g, CAGL0G02739g*, *CAGL0L03157g* and *CAGL0J04400g* are uncharacterized and annotated as the *Saccharomyces cerevisiae* orthologue of *RPA12*, *XBP1*, *DAL80*, and *HAP3*, respectively. Interestingly, three of the yeast orthologues (*RPA12*, *XBP1*, *DAL80*) have negative roles in transcription (Mai & Breeden, 1997; Marzluf, 1997; Yadav *et al*, 2016). Transcription regulatory network analysis by PathoYeastract (Monteiro *et al*., 2020) further showed that the orthologues of ∼35% macrophage infection-induced genes (n = 375 out of 1,078, respectively; Figure 2-figure supplement 1A, Supplementary File 4) are targets of Xbp1 in *S. cerevisiae* including a significant number of TF genes (n = 14; Figure 2-figure supplement 1B). In contrast, a much smaller set of orthologous genes (∼ 7%, n = 72, Figure 2-figure supplement 1A) is annotated as being *S. cerevisiae* Hap3 targets, while no information was available on the PathoYeastract database (Monteiro *et al*., 2020) for the other two repressors. These results suggest that the chronological transcriptional response upon macrophage phagocytosis involves the interplays between transcriptional repressors and activators and that the protein encoded by *CAGL0G02739g* (hereafter referred to as *CgXBP1*) likely plays a central role in orchestrating the overall response.

### CgXbp1 is crucial for the chronological transcriptional response during macrophage infection

We next deleted the *CgXBP1* gene and analysed the transcriptional response of the *Cgxbp1*Δ mutant to macrophages. RNAPII ChIP-seq time course analysis showed that a similar number of genes were transcribed in the mutant during macrophage infection (1,471 versus 1,589 genes in *Cgxbp1*Δ and wildtype, respectively) (Supplementary File 5) and ∼90% of the transcribed genes are common between wildtype and the mutant (Figure 2-figure supplement 1C), suggesting that CgXbp1 has little effect on the overall set of genes transcribed during macrophage infection. Notably, the *Cgxbp1*Δ mutant had a significantly higher number of genes activated at the earliest infection time point (0.5 h, Figure 2B, Supplementary File 5) as compared to wildtype (Figure 1C); e.g., 369 genes showed accelerated expression in the *Cgxbp1*Δ mutant, while 162 and 109 genes had an unchanged or delayed gene expression profile (Figure 2C).

Systematic GO analysis revealed multiple biological processes enriched among the genes with precocious activation in the *Cgxbp1*Δ mutant (0-0.5 h). They include processes like energy generation, chromatin assembly, cellular respiration and metabolism pathways such as TCA cycle, acetate catabolism, and amino acid, carboxylic acid, nucleotide and trehalose biosynthesis (Figure 3-figure supplement 1, Supplementary File 6). On the other hand, cell adhesion, host response and biofilm formation genes, which were up-regulated in wildtype cells during the late infection stage did not happen in the *Cgxbp1*Δ mutant within the 8 h infection duration examined (Figure 3-figure supplement 1).

Given that remodelling of carbon and nitrogen metabolism is crucial for the survival of fungal pathogens inside phagocytic cells (Lorenz and Fink, 2001; Rubin-Bejerano *et al*., 2003; Rai *et al*., 2012; Seider *et al*., 2014), we closely examined the expression patterns of TCA cycle and amino acid biosynthesis genes in wildtype and the *Cgxbp1*Δ mutant during macrophage infection. In wildtype cells, most genes of these two metabolic pathways were temporally induced with the maximal induction at 2h (Figures 3A&B). By contrast, the induction of these genes was advanced to 0.5 h (Figures 3A&B) and their overall expression levels were significantly higher (1.5 to 14.6 folds) in the mutant compared to wildtype, suggesting that CgXbp1 negatively regulates their expression (i.e., acting as a repressor) upon macrophage infection. Overall, the above results demonstrate that CgXbp1 is critical for the chronological transcriptional response of *C. glabrata* during macrophage infection.

**Figure 3:**
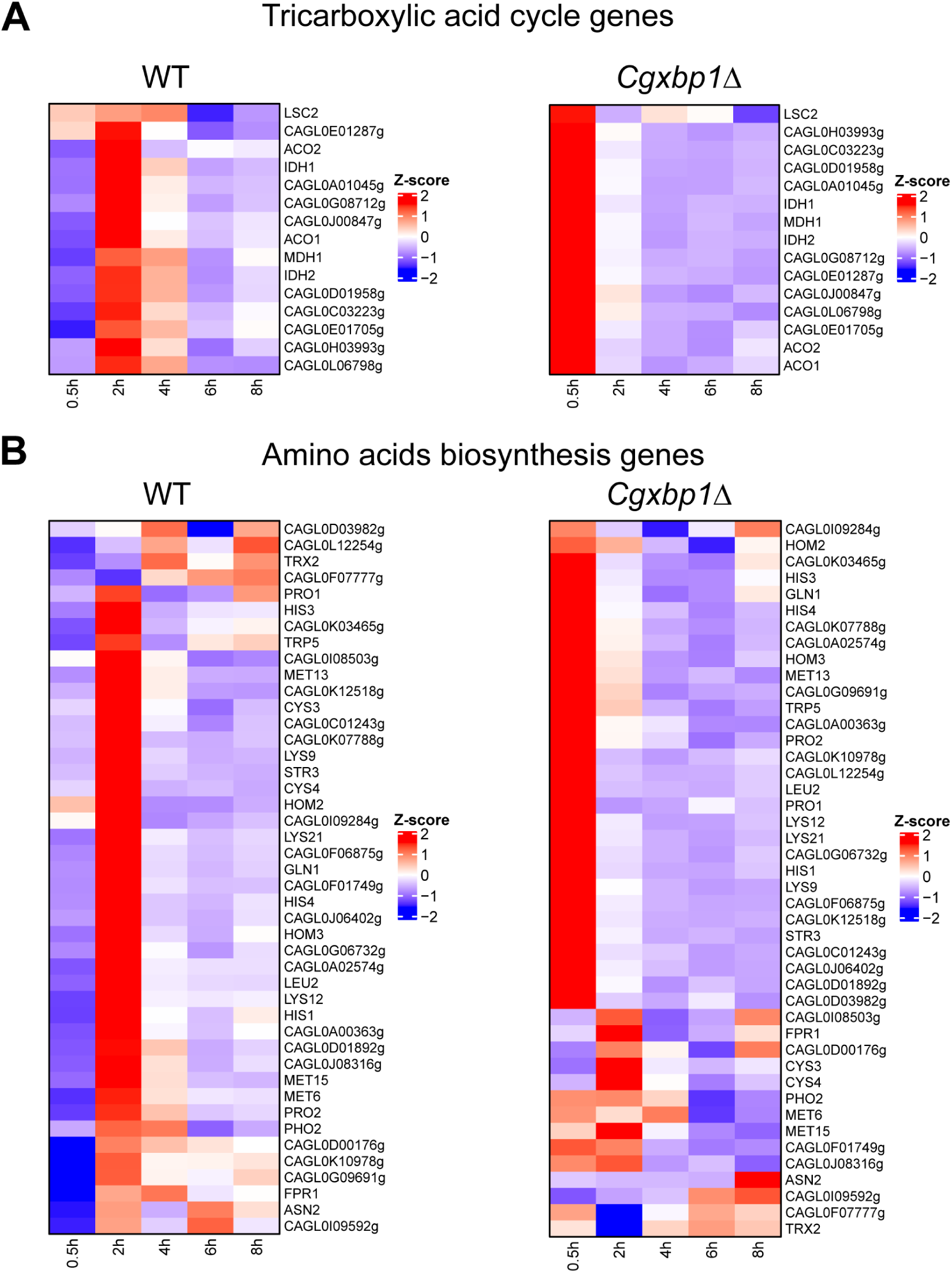
Loss of *CgXBP1* affects the expression level and timing of multiple genes of diverse physiological pathways upon macrophage phagocytosis. **(A & B)** Heat maps showing transcription activities of genes belonging to (A) TCA and (B) amino acid biosynthesis during THP1 macrophage infection in wildtype and the *Cgxbp1*Δ mutant.

### Loss of *CgXBP1* diminishes *C. glabrata* proliferation in human macrophages and attenuates virulence in the *Galleria mellonella* model of candidiasis

To examine if the altered transcriptional response in the *Cgxbp1*Δ mutant affects the survival of *C. glabrata* cells in macrophages, we compared the ability of wildtype and the *Cgxbp1*Δ mutant to survive in THP-1 macrophages. PMA-differentiated THP-1 macrophages were infected by wildtype and *Cgxbp1*Δ cells, and colony forming unit (CFU) assay was performed to determine the number of viable phagocytosed *C. glabrata* cells at 2, 8, and 24 hours post macrophage infection. No significant difference in CFUs between wildtype and *Cgxbp1*Δ cells was observed at 2 h (Figure 4-figure supplement 1), suggesting similar phagocytosis efficiency of THP-1 macrophages for the two strains. At 8 and 24 h post-infection, wildtype cells exhibited ∼ 1.6 and 5.1-fold increase in CFUs compared to that at 2h. Although the *Cgxbp1*Δ mutant was able to proliferate inside macrophages, it displayed significantly lower CFUs (∼20%) at both time points (1.3 and 3.9-fold) (Figure 4A). The reductions were rescued in the *Cgxbp1*-*pXBP1* complemented strain (Figure 4A). These results indicate that CgXbp1 is important for *C. glabrata* proliferation within macrophages.

**Figure 4:**
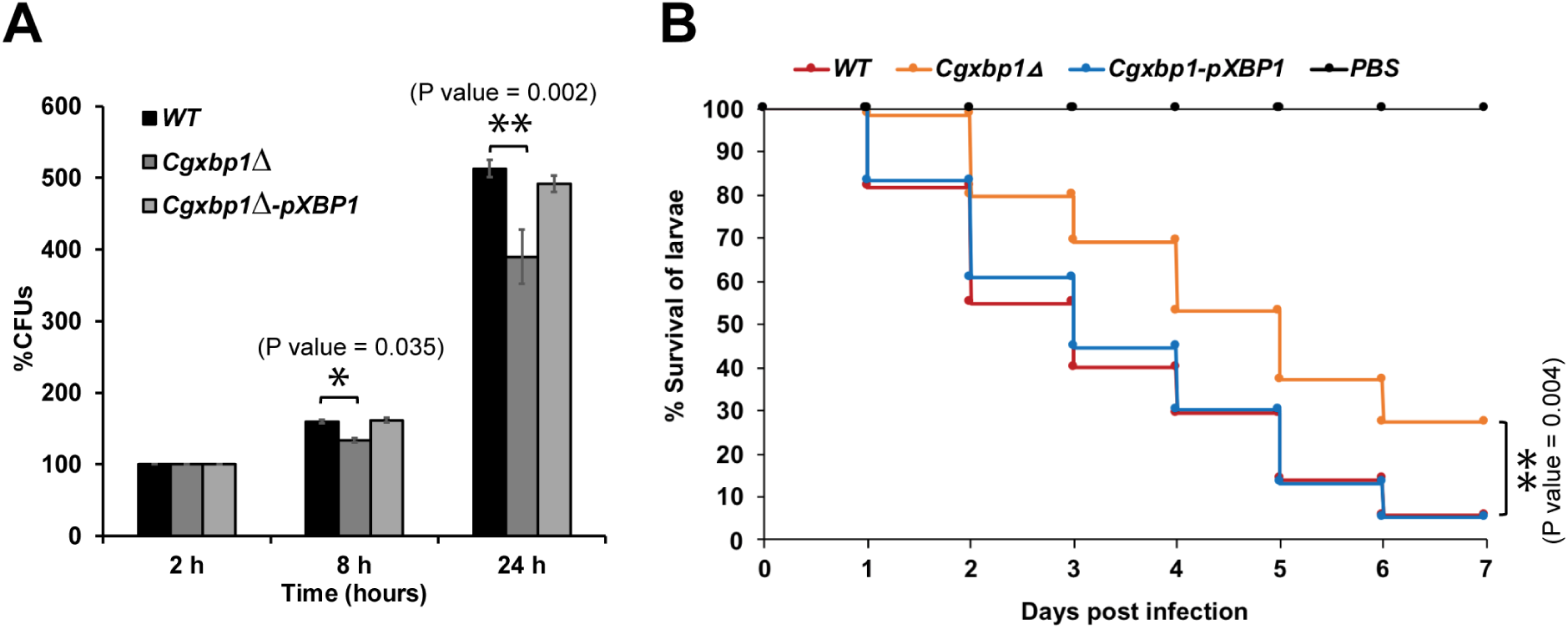
Loss of *CgXBP1* affects *C. glabrata* proliferation in human macrophages and attenuates virulence in the *Galleria mellonella* model of candidiasis. **(A)** Bar chart of CFUs obtained from *C. glabrata* cells harvested from THP-1 macrophages at indicated time points. Error bars represent the standard error of the mean (±SEM) from 3 independent experiments. Statistical significance was determined by two-sided unpaired Student’s *t*-test, *P value ≤ 0.05, **P value ≤ 0.01. **(B)** Cumulative survival curve of *G. mellonella* larvae infected with indicated *C. glabrata* strains. At least 16 larvae were used in each of three independent infection experiments. The graph represents the per cent survival of larvae infected with the indicated strains from three independent infection experiments. Statistical significance was determined by a two-sided unpaired Student’s *t*-test, **P value ≤ 0.01.

We next examined the virulence of the wildtype and *Cgxbp1*Δ strains using the *Galleria mellonella* model of *Candida* infection (Jacobsen, 2014). We infected *G. mellonella* larvae with the wildtype, *Cgxbp1*Δ, and complemented strains, and monitored the morbidity and mortality of infected larvae over seven days. Although worms injected with wildtype or *Cgxbp1Δ C. glabrata* cells (but not phosphate buffered saline [PBS]) turned dark grey within 4-6 h of infection due to melanin formation, which is a moth response to *C. glabrata* infection, and eventually died (Figure 4B), larvae injected with *Cgxbp1*Δ cells have a consistently slower mortality rate by ∼20-30% compared to larvae infected by wildtype cells, suggesting that the loss of Xbp1 function attenuated the virulence (Figure 4B). The attenuated virulence was rescued in the complemented strain (Figure 4B). Therefore, CgXbp1 is important for the survival of *C. glabrata* in human macrophages and virulence in the *in vivo* infection model.

### CgXbp1 targets the hierarchy of gene regulatory networks of diverse biological processes

The importance of CgXbp1 towards *C. glabrata*’s response during macrophage and *in vivo* infection models prompted us to characterize its direct genome-wide roles in more details. We attempted ChIP-seq analysis against CgXbp1^MYC^ during macrophage infection but failed to obtain consistent high-quality binding profiles from biological repeats (Pearson correlation *r* = 0.63), presumably due to dynamic responses of *C. glabrata* within the ever-changing macrophage microenvironment during macrophage infection. We, therefore, turned to a defined laboratory condition for the characterization. In *S. cerevisiae*, Xbp1 (ScXbp1) encodes a global transcription repressor that regulates 15% of genes during quiescency (Mai and Breeden, 1997; Miles *et al*., 2013). Pairwise amino acid sequence alignment showed reasonable conservation between ScXbp1 and CgXbp1, especially at the DNA binding domain and a few small scattered regions (Figure 5-figure supplement 1). Western blot analysis demonstrated that CgXbp1^MYC^ protein expression was significantly induced during macrophage infection (Figure 5A) as well as after 3 and 4 days of growth during which cells have already entered the stationary phase (Figure 5B). The sequence conservation and expression pattern indicate that CgXbp1 also plays roles during quiescency in addition to macrophage response.

**Figure 5:**
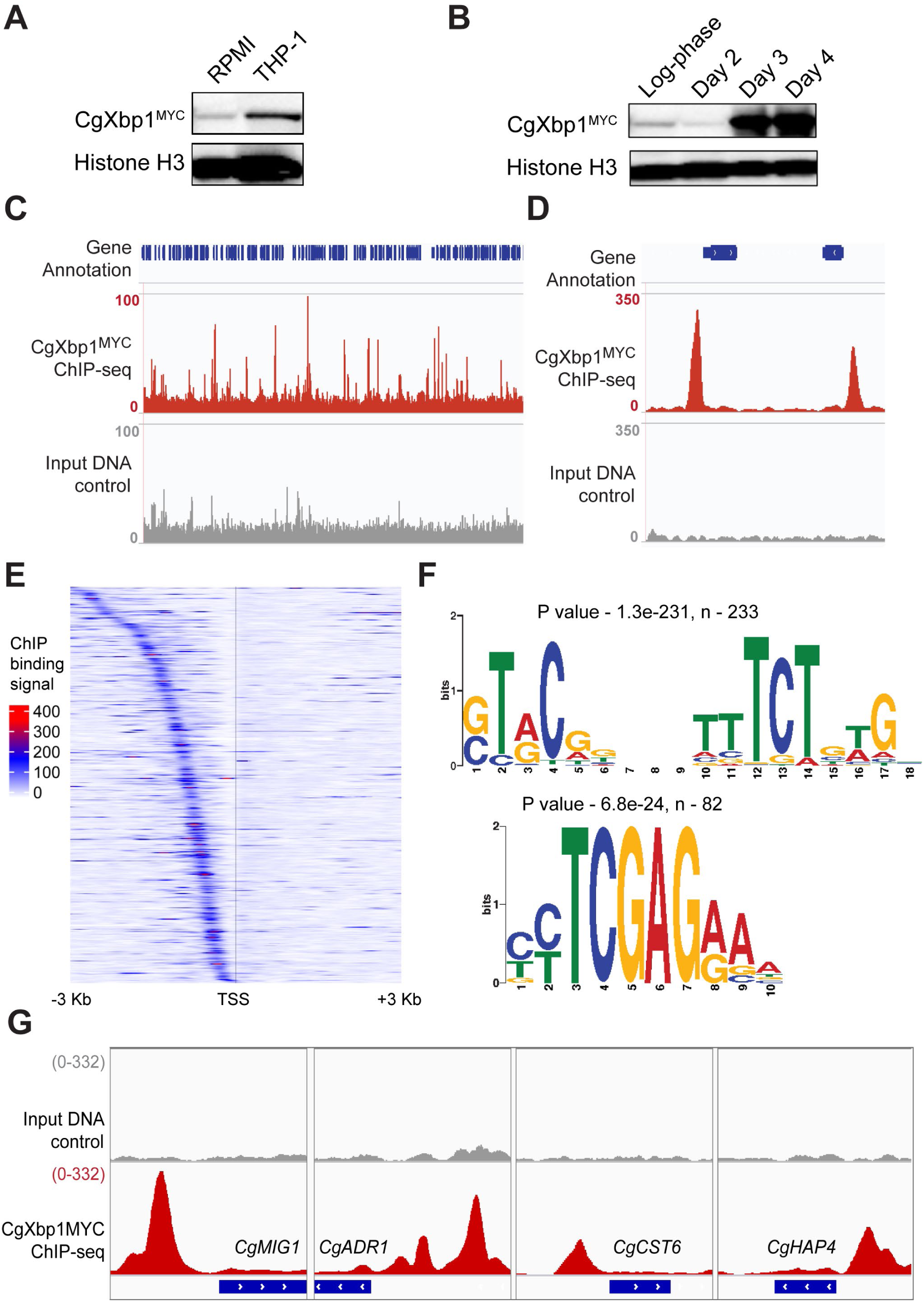
*CgXbp1* binds to a large number of genomic locations *in vivo*. **(A & B)** Western blot analysis of CgXbp1 expression (A) during THP1 macrophage infection and (B) at exponential (Log) and quiescence phases (Day 2, 3 and 4) (Figure 5-Source data). **(C & D)** Representative genome-browser screenshots showing CgXbp1^MYC^ ChIP-seq signal on (C) a chromosomal region and (D) the promoter of select genes. **(E)** A Heat map of ChIP-seq signal on promoters of CgXbp1 target genes. The colour scale indicates normalized ChIP-seq signal on 3 kb upstream and downstream flanking regions from the transcription start site (TSS) of the target genes. **(F)** Consensus DNA binding motifs enriched from CgXbp1 bound promoter sequences in CgXbp1 target genes by MEME analysis. **(G)** Genome-browser screenshots displaying CgXbp1 binding on *CgMIG1*, *CgADR1*, *CgCST6*, and *CgHAP4* promoters during the quiescence phase.

Next, we set out to identify CgXbp1^MYC^ genome-wide targets using ChIP-seq during the quiescence phase (day 4 culture). Unlike the attempt for macrophage infection, ChIP-seq against CgXbp1^MYC^ during quiescency produced highly correlated biological replicates (Pearson correlation *r* = 0.95) (Figure 5-figure supplement 2A&B). Distinct CgXbp1^MYC^ bindings were observed throughout the genome, while the input DNA control exhibited only background signals (Figures 5C&D). MACS2 analysis (Zhang *et al*., 2008; Feng *et al*., 2012) identified 519 CgXbp1^MYC^ binding sites with 420 located at the promoter (i.e., within 2 kb upstream of the translation start site) of 384 annotated genes (Figure 5E, Supplementary Files 7&8). Two over-represented DNA binding motifs (STVCN_7_TCT [where S represents G or C, V represents T or C] and TCGAG) were identified by *de novo* motif discovery using MEME suite (Bailey *et al*., 2009) (Figure 5F). While the TCGAG motif is similar to the consensus recognition sequence of *S. cerevisiae* Xbp1 ([TCGA], Mai & Breeden, 1997), the STVCN_7_TCT sequence appears to be composed of two halves with the second half showing some resemblance to the DNA binding motif of *S. cerevisiae* Azf1 (Figure 5-figure supplement 3A&B). Interestingly, the two motifs have different occurrence among the target promoters bound by CgXbp1^MYC^ with the STVCN_7_TCT motif occurring approximately three times more frequent than the TCGAG sequence (e.g., n = 233 versus n = 82, respectively) (Figure 5F, Supplementary File 9).

The direct functions of CgXbp1 were determined using GO analysis and were significantly associated with major biological processes important for host infection such as “transport”, “response to stress”, “carbon and nitrogen metabolism”, and “biofilm formation” (Table 2, Supplementary File 10). These functions are consistent with the above findings that CgXbp1 is important for *C. glabrata* response and survival in macrophages. More importantly, CgXbp1^MYC^ bindings were found at ∼10% of transcription regulators (n = 57 out of 623 genes annotated as transcription regulators), including eight regulators of various stresses (e.g., oxidative stress, osmotic stress, nutrient, pH, acetate, chemical and salts) and ten TF genes involved in carbon catabolite regulation (Figure 5G, Supplementary File 11) Therefore, CgXbp1 exerts its controls over key biological processes through regulating the hierarchy of gene regulatory networks.

**Table 2:** Table of significantly enriched and non-redundant GO-terms for biological processes among CgXbp1 target genes during quiescence phase, grouped based on biological functions.

|  | GO Term | P value | Genes in the background | CgXbp1 bound genes |
| --- | --- | --- | --- | --- |
| **Transport** | transmembrane transport | 1.19E-06 | 320 | 47 |
|  | carboxylic acid transport | 3.13E-04 | 69 | 14 |
|  | acetate transport | 1.40E-03 | 4 | 3 |
|  | xenobiotic transport | 2.48E-03 | 9 | 4 |
|  | oxygen transport | 5.16E-03 | 2 | 2 |
|  | fluconazole transport | 5.16E-03 | 2 | 2 |
|  | drug transport | 8.20E-03 | 12 | 4 |
|  | ammonium transport | 1.04E-02 | 7 | 3 |
|  | amino acid transport | 1.19E-02 | 44 | 8 |
|  | glucose transmembrane transport | 1.57E-02 | 8 | 3 |
| **Response to stress** | response to nutrient | 6.93E-04 | 58 | 12 |
|  | cellular response to chemical stress | 1.36E-03 | 184 | 25 |
|  | response to nitrosative stress | 1.46E-03 | 8 | 4 |
|  | response to acetate | 3.31E-03 | 5 | 3 |
|  | cellular response to oxidative stress | 5.09E-03 | 130 | 18 |
|  | response to osmotic stress | 1.07E-02 | 99 | 14 |
|  | response to oxidative stress | 1.13E-02 | 151 | 19 |
|  | response to salt stress | 1.70E-02 | 38 | 7 |
|  | response to pH | 1.76E-02 | 85 | 12 |
|  | response to acidic pH | 4.99E-02 | 12 | 3 |
| **Metabolism** | carbohydrate metabolic process | 1.50E-05 | 184 | 30 |
|  | carbohydrate biosynthetic process | 2.16E-04 | 59 | 13 |
|  | glucan biosynthetic process | 4.38E-04 | 27 | 8 |
|  | carbon catabolite regulation of transcription | 4.89E-04 | 41 | 10 |
|  | glutamine family amino acid biosynthetic process | 6.23E-04 | 22 | 7 |
|  | nitrogen utilization | 9.73E-04 | 12 | 5 |
|  | regulation of glycolytic process | 1.47E-02 | 3 | 2 |
|  | tricarboxylic acid cycle | 3.51E-02 | 26 | 5 |
| **Biofilm** | adhesion of symbiont to host | 4.38E-04 | 27 | 8 |
|  | fungal-type cell wall organization or biogenesis | 7.39E-04 | 227 | 30 |
|  | positive regulation of cell growth | 1.12E-03 | 24 | 7 |
|  | biofilm formation | 1.67E-03 | 81 | 14 |
|  | biological adhesion | 1.06E-02 | 70 | 11 |

### CgXbp1 is a negative regulator of fluconazole resistance

Several CgXbp1^MYC^ bound genes (*CgPDH1*, *CgPDR13*, *CgQDR2*, *CgAQR1* and *CgTPO1*; Figure 6A, Supplementary File 10) are important for *C. glabrata* drug resistance (Hallstrom *et al*., 1998; Miyazaki *et al*., 1998; Costa *et al*., 2016, 2013; Pais *et al*., 2016). Therefore, we examined whether CgXbp1 affects the resistance of *C. glabrata* to the antifungal fluconazole. Serial dilution spotting assay on solid media showed that *Cgxbp1*Δ mutant had higher resistance to fluconazole compared to wildtype (Figure 6B), and the resistance was restored to the wildtype level in the complemented strain (Figure 6B). Importantly, the altered resistance is not due to an intrinsic difference in growth rate between the two strains, as demonstrated by their indistinguishable growth rates in the absence of drug in liquid media (Figure 6C). However, in presence of fluconazole (64 µg/mL), the *Cgxbp1*Δ mutant was able to grow faster and to a higher density than wildtype (Figure 6C). It is interesting to note that a biphasic growth curve was observed for both strains in the presence of fluconazole, suggesting the existence of two populations of cells (sensitive versus resistant). Consistently, we also noted from the spotting assay relatively more resistant colonies in the *Cgxbp1*Δ mutant as compared to wildtype. To confirm this, we performed a CFU assay by plating an equal number of exponentially growing wildtype, *Cgxbp1*Δ mutant and complemented cells on YPD medium with or without fluconazole (64 µg/mL). The *Cgxbp1*Δ mutant displayed ∼8-fold higher CFUs on fluconazole compared to that of wildtype and the complemented strain (Figure 6D), demonstrating the loss of CgXbp1 function led to a larger population of resistant cells. In conclusion, these results demonstrate that CgXbp1 is a negative regulator of fluconazole resistance in *C. glabrata*.

**Figure 6:**
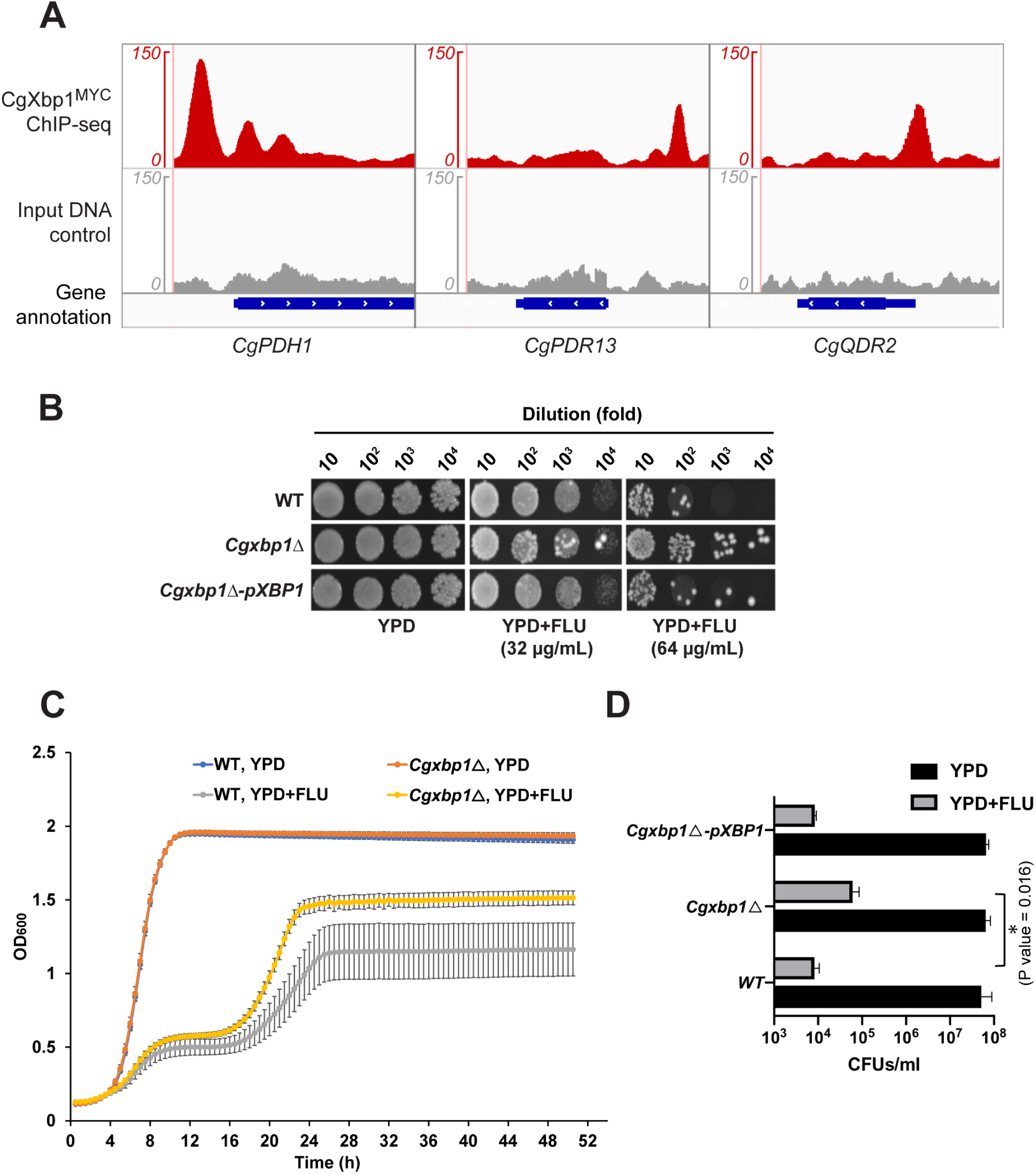
CgXbp1 regulates fluconazole resistance in *C. glabrata*. **(A)** Genome-browser screenshots showing CgXbp1^MYC^ binding on the promoter of genes encoding drug efflux pumps, *CgPDH1*, *CgPDR13*, and *CgQDR2.* **(B)** Serial dilution spotting assay on YPD medium in the presence of different fluconazole concentrations (0, 32 or 64 µg/mL). **(C)** Growth curve of wildtype and *Cgxbp1*Δ mutant in YPD medium in the presence or absence of fluconazole (64 µg/mL). Error bars represent mean ± SEM from four independent experiments. **(D)** Bar graph displaying CFUs/mL obtained for indicated strains on YPD agar plates in the presence or absence of fluconazole (64 µg/mL) post 3 days of spread plating. Error bars represent mean ± SEM from three independent experiments. Statistical significance was determined by a two-sided unpaired Student’s *t*-test, *P value ≤ 0.05.

## Discussion

*C. glabrata* is well known for its abilities to withstand azole antifungal drugs and to survive and grow inside phagocytic immune cells (Kaur *et al*., 2005; Rai *et al*., 2012). Through mapping genome-wide RNAPII occupancy, this work delineates the temporal transcriptional response of *C. glabrata* upon entry into macrophages, offering insights into the events occurring at different stages of macrophage infection. Our result reveals that ∼30% of *C. glabrata* genes are transcribed during the adaptation, survival and growth inside the alien macrophage microenvironments. At the most immediate response (0-0.5 h), *C. glabrata* activates adherence related genes to initiates adhesion to host surfaces. Concurrently, *C. glabrata* elicits specific responses to the nutrient-limiting microenvironment inside macrophages. This immediate response was followed by *C. glabrata* efforts to deal with oxidative and DNA-damage stresses (0-2 h), and at this time the phagocytosed *C. glabrata* cells were arrested at the G1-S phase of the cell cycle (Figure 1-figure supplement 2B). Subsequently (2-4 h), *C. glabrata* undergoes transcriptional remodeling to adjust its carbon metabolism, presumably to generate energy for future challenges, growth and/or proliferation. Consistently, processes necessary for growth such as global transcription, ribosome biogenesis and copper and iron ion homeostasis were activated around this time (2-6 h). These transcriptional activities indicate that *C. glabrata* are well-adapted to the macrophage microenvironment at this stage of infection. Lastly (8 h), *C. glabrata* induced genes associated with cell proliferation and biofilm formation, implying that they have overcome macrophage attacks and are ready to grow and divide.

It is noteworthy that despite sensing starvation within the first 0.5 h upon macrophage engulfment, *C. glabrata* does not activate alternate carbon catabolic pathways until 2 h. In addition, gene expression and translation-related genes show the lowest transcription levels (i.e., RNAPII occupancy) at this immediate stage (0.5 h) relative to the other time points (Group 6 genes in Figures 1C&D), indicating global suppression of gene expression in *C. glabrata* upon macrophage phagocytosis. A recent study showed that the fungal pathogen *Cryptococcus neoformans* also down-regulate translation during exposure to oxidative stress and suggested that translation suppression may facilitate the degradation of irrelevant transcripts during stress to facilitate proper gene expression and survival (Leipheimer *et al*., 2019). A similar strategy may be employed by *C. glabrata* in response to the stresses experienced upon macrophage phagocytosis. Another non-mutually exclusive possibility is that the suppression of gene expression helps *C. glabrata* to reserve energy and resources for coping with the hostile, nutrient-limiting macrophage environment.

Transcriptional responses are determined by the overall TFs activities in a cell. The RNAPII profiles revealed a panel of 53 TF genes expressed during macrophage infection. In particular, 39 of them were temporally induced and are promising candidate regulators for the stage-specific transcription responses. Of note, several uncharacterized TFs (CgXbp1, CgDal80 and CgRpa12) were strongly activated at the earliest infection stage. In *S. cerevisiae*, the orthologues of these TFs play negative regulatory roles (i.e., transcriptional repressors (Mai and Breeden, 1997; Marzluf, 1997; Yadav *et al*., 2016)), suggesting the importance of transcriptional repression in shaping the overall transcriptional response to macrophages (Figure 7). This was confirmed by the precocious transcriptional activation of a large number of genes in the *Cgxbp1*Δ mutant during macrophage infection. In addition, the ChIP-seq experiment revealed that CgXbp1 directly binds to the promoter of many TFs including 10 carbon catabolite regulators (Figure 5G, Supplementary File 11), suggesting that CgXbp1 indirectly represses the activation of many gene regulatory networks. This probably explains the delayed activation of the carbon catabolic pathway genes. Therefore, CgXbp1 exerts two-fold regulation; directly controlling downstream effector genes and indirectly affecting gene networks of diverse pathways via their hierarchies (i.e., TFs). Our overall findings suggest a regulatory model in which global transcriptional repression is established at the early infection stage to withhold transcriptional activation of certain genes whose functions are only required at later stages (Figure 7).

**Figure 7:**
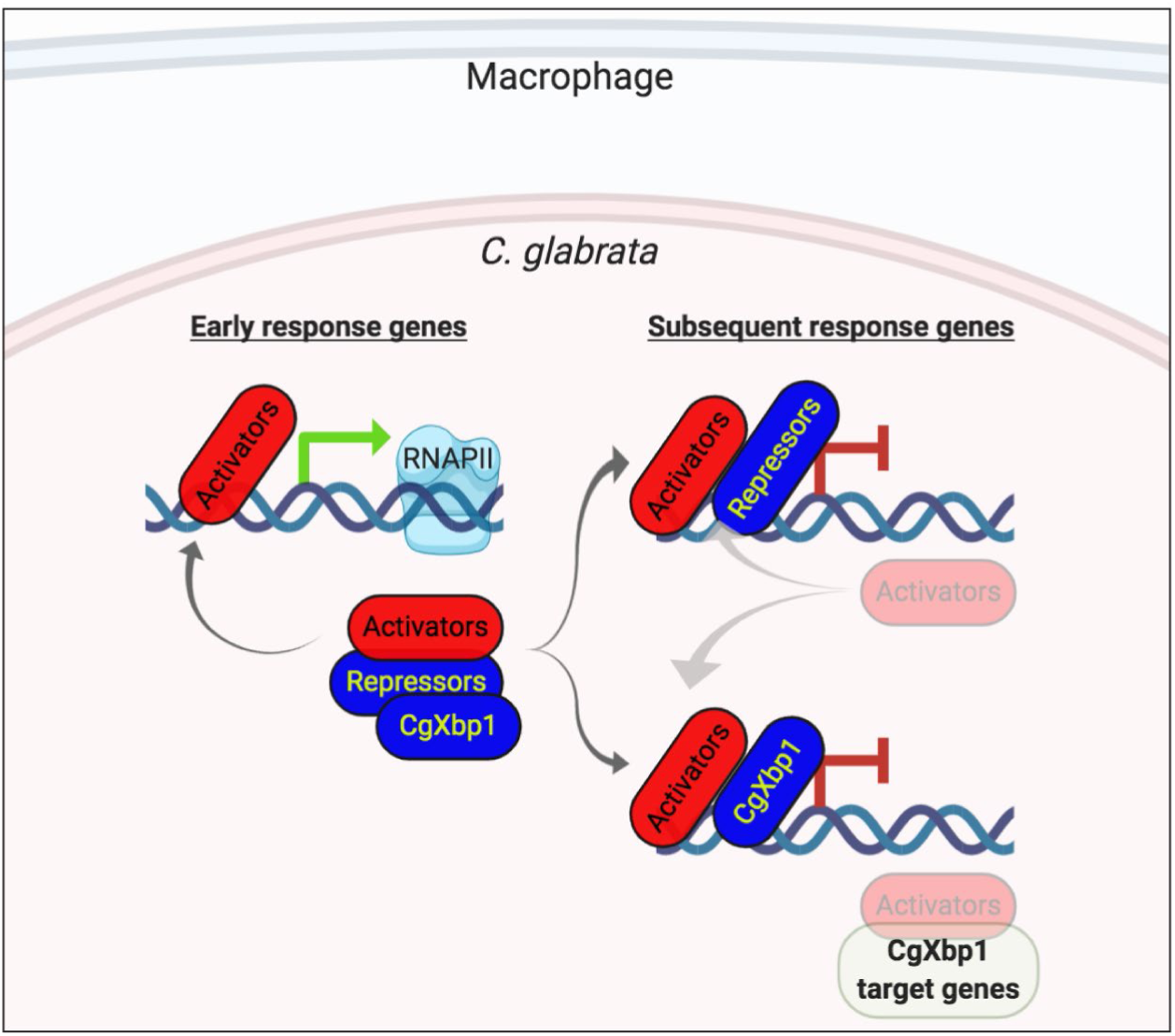
A schematic diagram illustrating a proposed model for the regulation of transcriptional responses in *C. glabrata* during macrophage infection. A model for the regulation of the chronological transcriptional responses of *C. glabrata* during macrophage infection. The model was created in BioRender.com.

It is noteworthy that the *S. cerevisiae* orthologues of the other two uncharacterized proteins CgDal80 and CgRpa12 negatively regulate genes involved in nitrogen and lipid metabolism, respectively (Hofman-Bang, 1999; Yadav *et al*., 2016). We postulate that these repressors also act in the same fashion as CgXbp1 to delay induction of other groups of genes (such as nitrogen and lipid metabolism genes) that are not necessary for the immediate response but are required at later stages of macrophage infection and/or for proliferation. Repression may be relieved by protein turnover of the repressors and/or through enhanced activation by transcriptional activators. Therefore, the interplays between transcriptional activators and repressors (Figure 7) are crucial in shaping the dynamic transcriptional response during macrophage infection.

An attempt to identify CgXbp1 genome-wide binding targets during macrophage infection was made, however, we only managed to obtain one good quality dataset. Nevertheless, analysis of the CgXbp1 ChIP-seq result during quiescence (4-day-old culture) showed that CgXbp1 targets ∼10% of all *C. glabrata* genes. Among the CgXbp1 binding sites, the motifs TCGAG and STVCN_7_TCT were significantly enriched. The former motif is similar to the recognition motif of ScXbp1 (TCGA), which was identified through promoter analysis and *in vitro* gel retardation and footprint experiments using recombinant ScXbp1 (Mai and Breeden, 1997; Miles *et al*., 2013), and is likely bound by CgXbp1 as a monomer or homo-dimer. Interestingly, the latter motif (STVCN_7_TCT) was found at a higher frequency (∼3 fold) than the common TCGAG motif from the CgXbp1^MYC^ binding sites, suggesting that CgXbp1 can also form a dimer with another transcription factor that recognizes the STVCN_7_TCT sequence and that this hetero-dimer controls a larger number of genes than by CgXbp1 alone.

The CgXbp1^MYC^ ChIP-seq data also confirms the transcriptional phenotype of the *Cgxbp1*Δ mutant demonstrating that CgXbp1 is a pivotal regulator of many processes that contributes to the survival inside phagocytic cells (Lorenz and Fink, 2001; Rubin-Bejerano *et al*., 2003; Rai *et al*., 2012; Seider *et al*., 2014). In addition, the data further revealed CgXbp1 as an important regulator of drug resistance, linking *C. glabrata* macrophage response and drug resistance. Therefore, this work uncovers a multifaceted transcriptional regulator important for the dynamic responses during macrophage infection and antifungal drug resistance in the human fungal pathogen *C. glabrata*.

## Methods

### Culture conditions for *C. glabrata* and THP-1 macrophages

*C. glabrata* strain BG2 was used as the wildtype in all experiments. A single colony of *C. glabrata* strains were cultured overnight (14-16 h) in YPD medium at 30°C and 200 rpm in a shaker incubator. To obtain quiescence *C. glabrata* cells, cultures were grown in YPD medium at 30°C and 200 rpm for four days. The THP-1 cell line was obtained from ATCC (TIB 202). THP-1 cells were grown in RPMI medium supplemented with 20 mM glutamine, antibiotic (penicillin-streptomycin, 1X), and 10% heat-denatured serum at 37°C with 5% CO_2_ in a cell culture incubator.

### *C. glabrata* infection of macrophages for RNAPII ChIP-seq

Macrophage infection assays were done as described previously (Rai *et al*., 2013). THP-1 monocytes were grown till 80% confluence, harvested, and re-suspended in RPMI medium at a cell density of 1 × 10^6^ cells/mL. For macrophage differentiation, Phorbol-12-myristate 13-acetate (PMA) was added to the THP-1 monocytes at a final concentration of 16 nM. Approximately 10 million cells were seeded in 100 mm culture dishes and incubated for 12 hours at 37°C with 5% CO_2_ in a cell culture incubator. Subsequently, the culture medium was replaced with fresh pre-warmed complete RPMI medium to remove PMA, and cells were allowed to recover in the absence of PMA for 12 hours. Macrophage differentiation and adherence were confirmed under the microscope. Overnight grown *C. glabrata* cells were harvested, washed with PBS and finally suspended in complete RPMI medium at a density of 10^8^ yeasts/ml. To infect THP-1 macrophages, 500 μL yeast cell suspension (5×10^7^ yeast cells) was added to each culture dish containing differentiated THP-1 macrophages at a MOI of 5:1. Post 0.5 h macrophage infection, THP-1 macrophages were crosslinked using formaldehyde at a final concentration of 1% for 20 minutes before 1.5 mL of 2.5 M glycine (a final concentration of 320 nM) was added to stop the crosslinking reaction. For the remaining time points (2 h, 4 h, 6 h and 8 h), culture dishes were washed gently with PBS three times to remove non-phagocytosed yeast cells, and the medium was replaced with fresh pre-warmed RPMI medium. The infected culture was further incubated until the indicated infection times before formaldehyde crosslinking as described above, infected macrophage cultures were harvested and washed three times with ice-cold TBS before storing in -80°C freezer till chromatin extraction.

### ChIP and Illumina sequencing library preparation

Chromatin was prepared using a previously described protocol (Fan et al., 2008) with modifications. Briefly, the infected macrophage cell pellet was resuspended in 400 µL FA lysis buffer and 10 µL of 100 mM PMSF solution in the presence of 100 µL equivalent zirconium beads and lysed using six 3-min cycles at maximum speed in a Bullet Blender^®^ (Next Advance) homogeniser with at least 3 min of cooling on ice in between each cycle. Cell lysate was transferred to a new 1.5 mL tube and centrifuged at 2500 g for 5 min in a microcentrifuge. The supernatant was discarded, and the resultant pellet was re-suspended in 500 µL FA lysis buffer, and then transferred to a 2 mL screw-cap tube. Sonication was carried to shear the crosslinked chromatin (cycles of 10 sec on and 15 sec off sonication for a total of 30 min sonication time), and chromatin was stored in the -80 °C freezer until use. Chromatin immuno-precipitation was carried out using 2 µL of a commercially available anti-RNA polymerase II subunit B1 phospho-CTD Ser-5 antibody (Millipore, clone 3E8, cat. no. 04-1572) for RNAP II, and anti-MYC tag antibody (Santa Cruz, cat. no. 9E10) for CgXbp1^MYC^. The sample was gently mixed on an end-to-end rotator at room temperature for 1.5 h, and 10 µL of packed protein A sepharose beads (GE Healthcare cat. no. 17-0618-01) were then added. The mixture was further incubated at room temperature for another 1.5 h with gentle mixing. Immuno-precipitated material was washed twice with FA lysis buffer (150 mM NaCl), and once with FA lysis buffer (500 mM NaCl), LiCl wash buffer and TE buffer before elution in 100 µL of elution buffer, as described previously(Wong and Struhl, 2011). Eluted DNA was decrosslinked at 65°C overnight and purified using the QIAGEN PCR purification kit (cat. no. 28104). Sequencing library was generated using a multiplex Illumina sequencing protocol (Wong *et al*., 2013) and sequenced using the Illumina HiSeq2500 platform at the Genomics and Single Cells Analysis Core facility at the University of Macau.

### Bioinformatics and ChIP-seq data analyses

Raw fastq sequences were quality-checked using FastQC (http://www.bioinformatics.babraham.ac.uk/projects/fastqc/) aligned against the *C. glabrata* reference genome (CBS138_s02-m07-r06) using bowtie2 (Langmead and Salzberg, 2012). To visualise the ChIP-seq data on the IGV (integrated genome viewer (Thorvaldsdóttir et al., 2012)), aligned reads were processed by MACS2 (Zhang *et al*., 2008), and BigWig files were generated using ‘bedSort’ and ‘bedGraphToBigWig’ commands from UCSC Kent utils (Kent *et al*., 2010). Samtools (version 1.9) was used to index the resultant BAM file and check for alignment statistics. For RNAPII ChIP-seq analysis, elongating RNAPII occupancy was measured by first counting the number of reads over the gene body for all annotated genes (n = 5,311) and then normalising to gene length and sequencing depth using an in-house Perl script, and was expressed as normalised RNAPII ChIP-seq read counts. RNAPII occupancies for all genes were ranked from high to low as shown in Figure 1-figure supplement 1, and manually inspected on the IGV to empirically determine a filtering cut-off that can reliably identify genes with significant and true RNAPII ChIP-seq signals. The normalised RNAPII ChIP-seq read counts values ≥ 12 and ≥ 25 were determined for wildtype and the *Cgxbp1*Δ mutant, respectively. These values are approximately three times higher than the ChIP-seq signal at background regions (3.2 and 7.0 for wildtype and the *Cgxbp1*Δ mutant, respectively). To ensure that lowly expressed but transcriptionally induced genes were not missed, we searched for genes with the high standard deviation among RNAPII binding signals across the five time points and empirically determined a cut-off (SD ≥ 2.25 and ≥ 4.00 for wildtype and the *Cgxbp1*Δ mutant, respectively) that includes most, if not all, genes with significant active transcription and/or changes in its level across the time course. This standard deviation-based approach identified 68 and 38 additional genes for wildtype and the *Cgxbp1*Δ mutant, respectively. The lists of shortlisted genes for wildtype and the *Cgxbp1*Δ strains are given in Supplementary Files 1 and 5, respectively. Heatmap, k-means clustering and correlation plots were generated using an online tool FungiExpresZ (https://cparsania.shinyapps.io/FungiExpresZ/). GO-term enrichment, and GO slim mapping analyses were performed on the *Candida* genome database (Skrzypek *et al*., 2017) (http://www.candidagenome.org/GOContents.shtml) and FungiDB (Stajich *et al*., 2012) (https://fungidb.org/fungidb/). Transcription regulatory networks between transcription factors and their target genes (Figure 2-figure supplement 1A&B) were generated by the ‘Rank by TF’ function of PathoYeastract (Monteiro *et al*., 2020) (http://pathoyeastract.org/cglabrata/formrankbytf.php) using published regulatory information of *S. cerevisiae*. For CgXbp1 ChIP-seq analysis, peak calling was done by MACS2 (Feng *et al*., 2012) using the parameters [macs2 pileup --extsize 200] and then normalized [macs2 bdgopt]. Peaks obtained from MACS2 were assigned to the nearest gene promoter using an in-house script. ChIP signal intensity at the 200 bp flanking regions of the peak summit from both replicates was used to determine correlation between the biological replicates (Figure 1-figure supplement 6, Figure 5-figure supplement 2).

### Generation of the CgXbp1^MYC^, *Cgxbp1*Δ and *Cgxbp1*Δ*-pXBP1* complemented strains

CgXbp1^MYC^ strain was generated as described previously (Qin *et al*., 2019). Briefly, a transformation construct was generated using 1kb of 5’ and 3’ fragments, flanking the stop codon of *CAGL0G02739g* gene, with a ‘MYC-*hph*’ cassette between the two fragments. The 5’ fragment for CgXbp1^MYC^ strain was amplified using primers 5’-ATATCGAATTCCTGCAGCCCTCCATGGTACATTGCAAAAC-3’, and 5’-TTAATTAACCCGGGGATCCGCACATTCTCTTGAAGATGGG-3’ from *CAGL0G02739g* gene, and 3’ fragment was same as for *Cgxbp1*Δ mutant. Hygromycin resistant yeast colonies were selected and tagging was confirmed PCR and Sanger sequencing. To create the *Cgxbp1*Δ mutant, 1 kb of 5’ and 3’ flanking regions of the *CAGL0G02739g* gene were amplified using PCR with the primers 5’-ATATCGAATTCCTGCAGCCCGGCCAACCCCACTTCGAGGA-3’ and 5’-TTAATTAACCCGGGGATCCGTTAGTGATTTTGTAGTATGG-3’ for the 5’ flanking region and 5’-GTTTAAACGAGCTCGAATTCTCAAACATAATATAGTCATC-3’ and 5’-CTAGAACTAGTGGATCCCCCGAGAAGTTTTGGGTTGTACG-3’ for the 3’ flanking region. A transformation construct was created as described previously (Qin et al., 2019) using the ‘*hph’* cassette encoding hygromycin resistance as the selectable marker and used to transform the *C. glabrata* wildtype strain (BG2). Hygromycin resistant yeast colonies were checked for deletion of the *CgXBP1* gene using PCR with the primers from gene internal regions (5’-TGGTGCTTTGGACGCTACAT-3’ & 5’-TCATCGCAAAAGCAATTGGACA-3’). To generate the complemented strain, *CgXBP1* ORF was first amplified using the forward (5’- GAATTCATGAGACTCACAGACTCGCCGCT-3’) and reverse (5’- GTCGACTTACACATTCTCTTGAAGATGGGT-3’) primers from *C. glabrata* genomic DNA, digested with *EcoR*I and *Sal*I, and cloned between *EcoR*I and *Sal*I restriction sites of a CEN/ARS episomal plasmid, pCN-PDC1. The resultant plasmid, pXBP1, carrying *CgXBP1* ORF was transformed into *Cgxbp1*Δ mutant, and the resultant transformed complemented strain was selected on YPD plates carrying NAT (100µg/mL).

### Cell cycle analysis of intracellular *C. glabrata* cells in THP-1 macrophages

For cell cycle analyses, THP-1 macrophages were infected with *C. glabrata* cells in 24-well cell culture plate as described above. In control wells, we inoculated an equal number of *C. glabrata* cells to RPMI medium. Post 2 h incubation, *C. glabrata* cells were harvested and washed twice with 1 mL PBS. Next, harvested cells were fixed by re-suspending them in 1 mL of 70% ethanol, followed by incubation at room temperature on a rotator for 60 minutes. Fixed cells were pelleted and re-suspended in 1 mL of 50 mM sodium citrate (pH 7.0), and were sonicated for 15 seconds at 30% power to re-suspend cell aggregates. Subsequently, samples were treated with RNase cocktail (0.3 µL, Ambion cat. no. AM2286) at 37°C for 1 h to remove RNA, washed with PBS, and stained with propidium iodide (PI) for 1 hour. Cells were then passed through a 40 μm membrane filter and were analysed on the BD Accuri C6 flow cytometer (excitation: 488nm Laser, filter: 585/40, and detector: FL2).

### *C. glabrata* infection of macrophages for determining viability using colony forming unit assay

Macrophage fungi infection assays were done as described earlier (Rai *et al*., 2013). To prepare macrophages for infection assay, THP-1 monocytes were grown till 80% confluence, harvested, and resuspended to a cell density of 10^6^ cells/ml in complete RPMI medium. Phorbol-13-myrstyl-acetate (PMA) was added to the cell suspension to 16 nM final concentration, mixed well, and 1 million cells were seeded in each well of a 24 well cell culture plate. Cells were incubated for 12 hours in a cell culture incubator, the medium was replaced with fresh pre-warmed complete RPMI medium, cells were allowed to recover from PMA stress for 12 hours. Macrophage differentiation and adherence were confirmed under the microscope. Overnight grown *C. glabrata* cells were harvested, washed with PBS and adjusted to 0.1 OD^600^ and resuspended in complete RPMI medium. 100 µL yeast cell suspension was added to each well of the 24 well culture plate containing differentiated THP-1 macrophages. Post 2-hour co-incubation, wells were washed three times with PBS to remove non-phagocytosed yeast cells and the medium was replaced. At indicated time post-infection, the supernatant was aspirated out from the wells, and macrophages were lysed in sterile water and incubated for 5 minutes for lysing the macrophages. Lysates containing fungal cells were collected, diluted appropriately in PBS, and plated on YPD mediums. Plates were incubated for two days at 30°C and colonies were counted after 48 hours. The viability of *C. glabrata* cells was determined by comparing the colony forming units.

### *Galleria mellonella* infection assay for virulence analyses

Indicated *C. glabrata* strains were grown in YPD medium overnight, washed with PBS thrice, and resuspended in PBS to a final cell density of 10^8^ cells/ml. Next, 20 µL of this cell suspension carrying 2 × 10^6^ *C. glabrata* cells were used to infect *G. mellonella* larvae. The infection was carried out three independent times, each on 16 to 20 larvae. An equal volume of PBS was injected into the control set of larvae. Infected larvae were transferred to a 37°C incubator, and monitored for melanin formation, morbidity and mortality for the next seven days at every 24 hours. The number of live and dead larvae was noted for seven days, and the percentage of *G. mellonella* larvae survival was calculated.

### Serial dilution spotting assay

*C. glabrata* strains were grown in YPD medium for 14-16 h at 30°C under continuous shaking at 200 rpm. Cells were harvested from 1 ml culture, washed with PBS, and were diluted to an OD_600_ of 1. Next, five ten-fold serial dilutions were prepared from an initial culture of 1 OD_600_. Subsequently, 3 μL of each dilution was spotted on YPD plates with or without fluconazole (32 & 64 μg/mL). Plates were incubated at 30°C and images were captured after 2-8 days of incubation.

### Growth curve analyses

A single colony of the indicated strains was inoculated to liquid YPD medium and grown for 14-16 h. The overnight grown culture was used to inoculate to YPD medium with or without 64 µg/mL fluconazole at an initial OD_600_ of 0.1 in a 96-well culture plate. The culture plate was transferred to a 96 well-plate reader, Cytation3, set at 30°C and 100 rpm. The absorbance of cultures was recorded at OD_600_ nm at regular intervals of 30 minutes over a period of 48 h. Absorbance values were used to plot the growth curve.

### Protein extraction and western blotting

For protein extraction from macrophage-internalized *C. glabrata* cells, THP-1 macrophages were infected as described above. At the indicated time post-infection, macrophages were lysed in sterile chilled water, and phagocytosed *C. glabrata* cells were recovered and washed with 1X TBS buffer, transferred into 1.5 ml microcentrifuge tubes, and stored at -80°C until use. *C. glabrata* cell pellets were resuspended in 1X lysis buffer (50 mM HEPES, pH 7.5; 200 mM NaOAc, pH 7.5; 1 mM EDTA, 1 mM EGTA, 5 mM MgOAc, 5% Glycerol, 0.25% NP-40, 3 mM DTT and 1 mM PMSF) supplemented with protease inhibitor cocktail (Roche). Zirconium beads equivalent to 100 µL volume was added in microcentrifuge tubes and resuspended cells were lysed by 6 rounds of bead beating on a bullet blender. The sample was centrifuged at 12,000g at 4°C for 10 minutes. Supernatant was carefully transferred to a new tube, and the resultant protein sample was quantified using Biorad protein assay kit (DC protein assay kit, cat. no. 5000116), and stored in -80°C freezer. For western analysis, 25 µg of protein samples were resolved on 12% SDS-PAGE gel and blotted on methanol activated PVDF membrane (350 mA, 75 minutes in cold room). PVDF membrane was transferred to 5% fat-free milk prepared in 1X TBST for blocking and incubated for 1 hour. Membranes were probed with appropriate primary (anti c-MYC antibody, Santa Cruz, cat. no. 9E10 and anti-Histone H3 antibody, Abcam, cat. no. ab1791) and secondary (goat anti-mouse IgG, Merck Millipore, cat. no. AP124P) antibodies, and Blots were developed by chemiluminescence based ECL western detection kit (GE Healthcare, cat. no. RPN2236) on Chemidoc™ gel imaging system.

## Data availability

RNAPII ChIP-seq and CgXbp1^MYC^ ChIP-seq data are available from the NCBI SRA database under the accession number PRJNA665114 and PRJNA743592, respectively.

## Acknowledgements

We thank members of the Wong laboratory for their valuable comments throughout the study. We acknowledge the services and technical supports from the Genomics and Single Cell Analysis Core and the Drug and Development Core of the Faculty of Health Sciences at the University of Macau. This work was performed in part at the High-Performance Computing Cluster (HPCC), which is supported by the Information and Communication Technology Office (ICTO) of the University of Macau. We thank Lakhansing Pardeshi and Zhengqiang Miao, for Bioinformatics supports and Jacky Chan for technical supports on the HPC. This work was supported by the Research Services and Knowledge Transfer Office of the University of Macau (Grant number: MYRG2019-00099-FHS) and the Collaborative Research Fund Equipment Grant (C5012-15E) from the Research Grant Council, Hong Kong Government.

## Author contributions

M.N.R. and K.H.W conceived the study, designed experiments and interpreted data. M.N.R. and R.R. performed experiments. M.N.R, C.P. and N.S. performed Bioinformatics analysis. K.T. performed Illumina sequencing and provided technical supports. K.T. and K.H.W. provided funding and resources. M.N.R. and K.H.W. wrote the manuscript.

---

**Figure 1-figure supplement 1:**
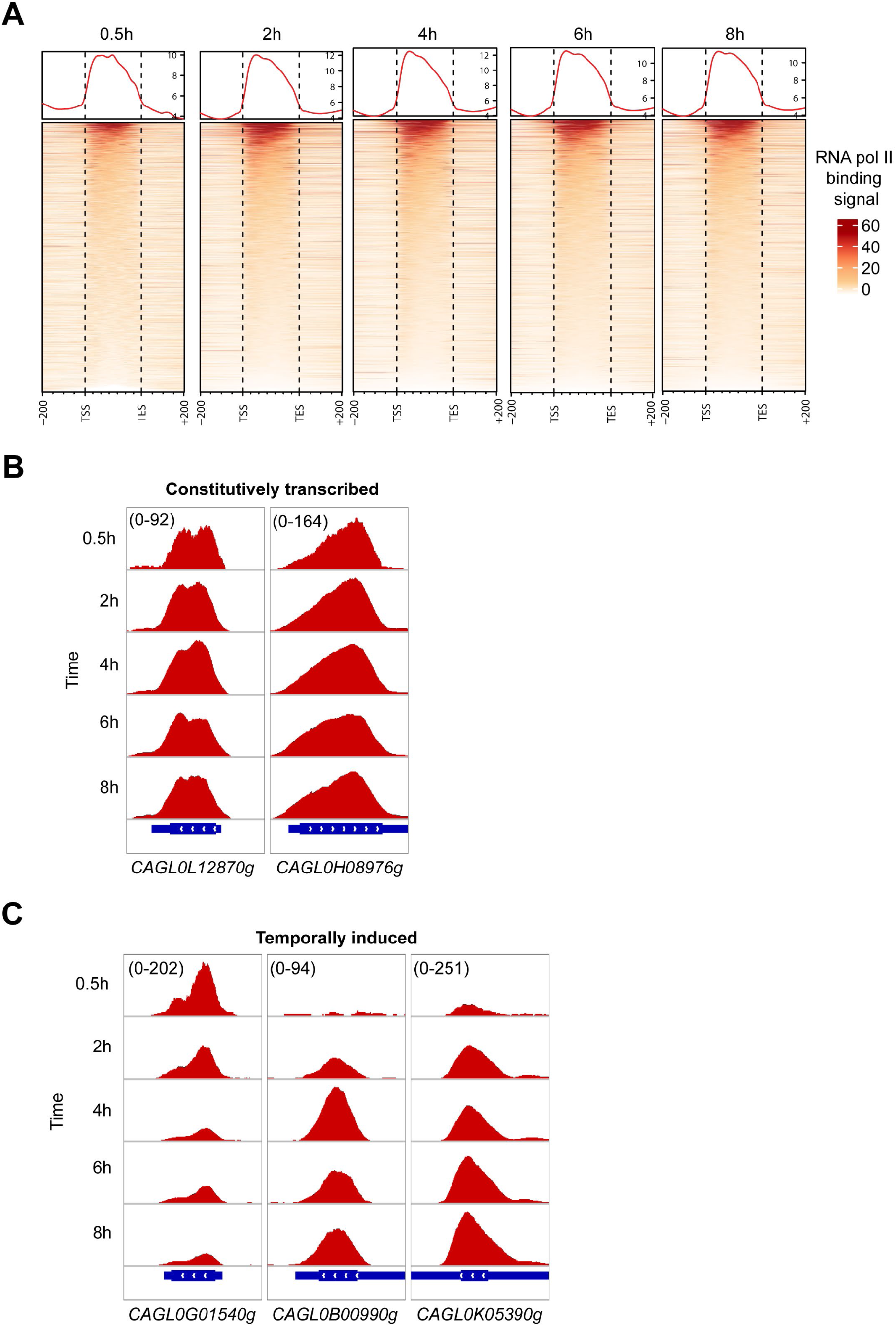
High-resolution RNAPII ChIP-seq can capture genome-wide active and temporally induced transcription activities in *C. glabrata* during macrophage infection. **(A)** A heatmap displaying RNAPII ChIP-seq signal over the gene body and 200 bp upstream and downstream regions for all *C. glabrata* genes after 0.5, 2, 4, 6 and 8 h of THP-1 macrophage infection. The colour scale represents the normalized RNAPII ChIP-seq signal. **(B)** Genome browser screenshots showing RNAPII ChIP-seq signal on selected constitutively transcribed genes. **(C)** Genome browser screenshots showing RNAPII ChIP-seq signal on selected temporally induced *C. glabrata* genes. Numbers in the square brackets indicate the y-axis scale range of normalized RNAPII ChIP-seq signal used for the indicated genes across different datasets.

**Figure 1-figure supplement 2:**
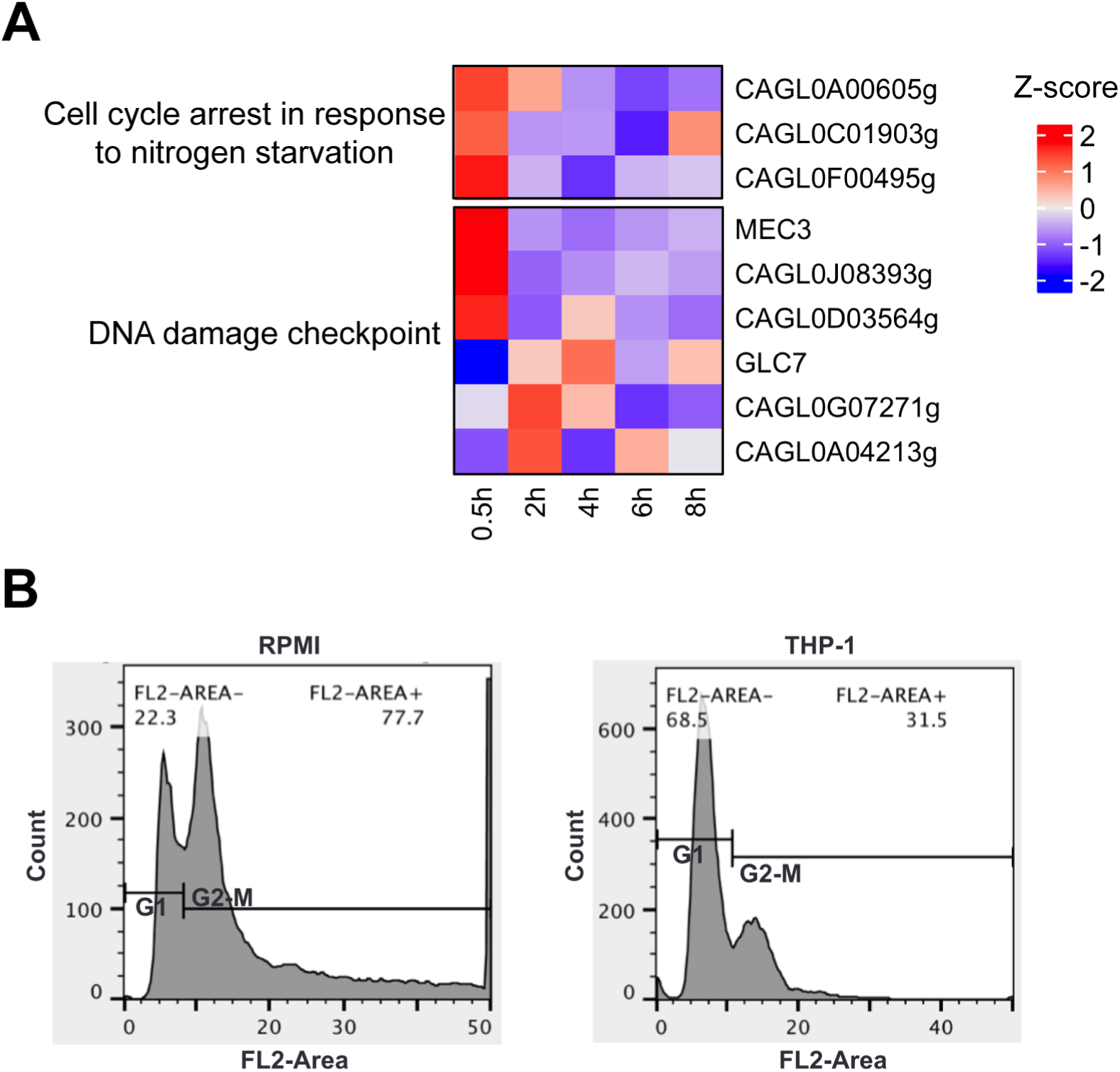
*C. glabrata* undergoes cell cycle arrest upon macrophage phagocytosis. **(A)** Heatmaps showing the expression pattern for cell cycle and DNA damage checkpoint genes during macrophage infection. The colour scale represents the Z-score of the normalized RNAPII ChIP-seq signal. **(B)** A density plot displaying the distribution of *C. glabrata* cells at different cell cycle stages based on FACS analysis at 2 h after THP-1 macrophage infection.

**Figure 1-figure supplement 3:**
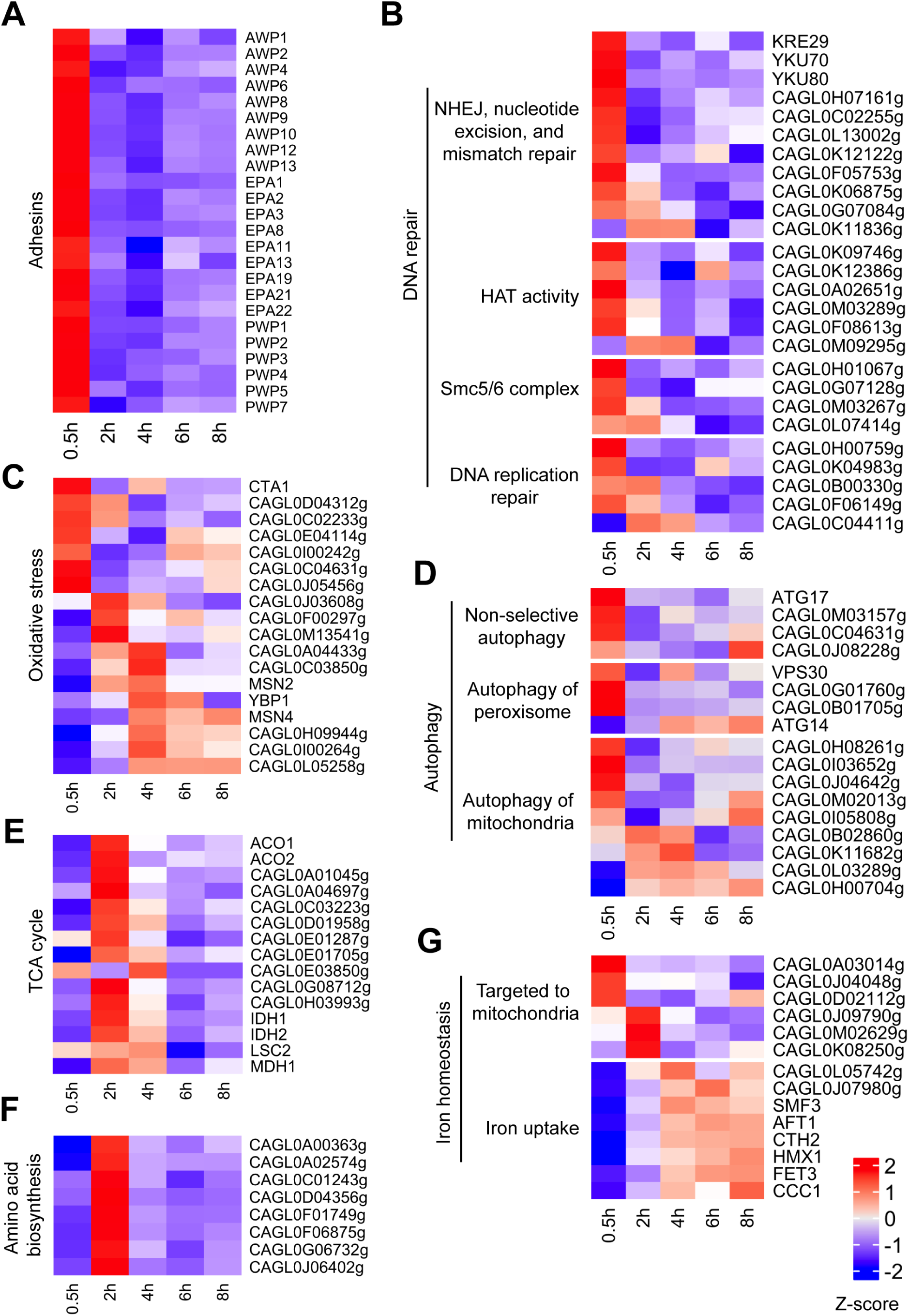
Virulence-centric biological processes are temporally activated in *C. glabrata* at different stages of macrophage infection. **(A-G)** Heatmaps displaying the expression pattern for genes associated with biological processes (A) Adhesion, (B) DNA repair, (C) Response to oxidative stress, (D) Autophagy, (E) TCA cycle, (F) Amino acid biosynthesis and (G) Iron homeostasis during THP-1 macrophage infection. The colour scale represents the Z-score of the normalized RNAPII ChIP-seq signal.

**Figure 1-figure supplement 4:**
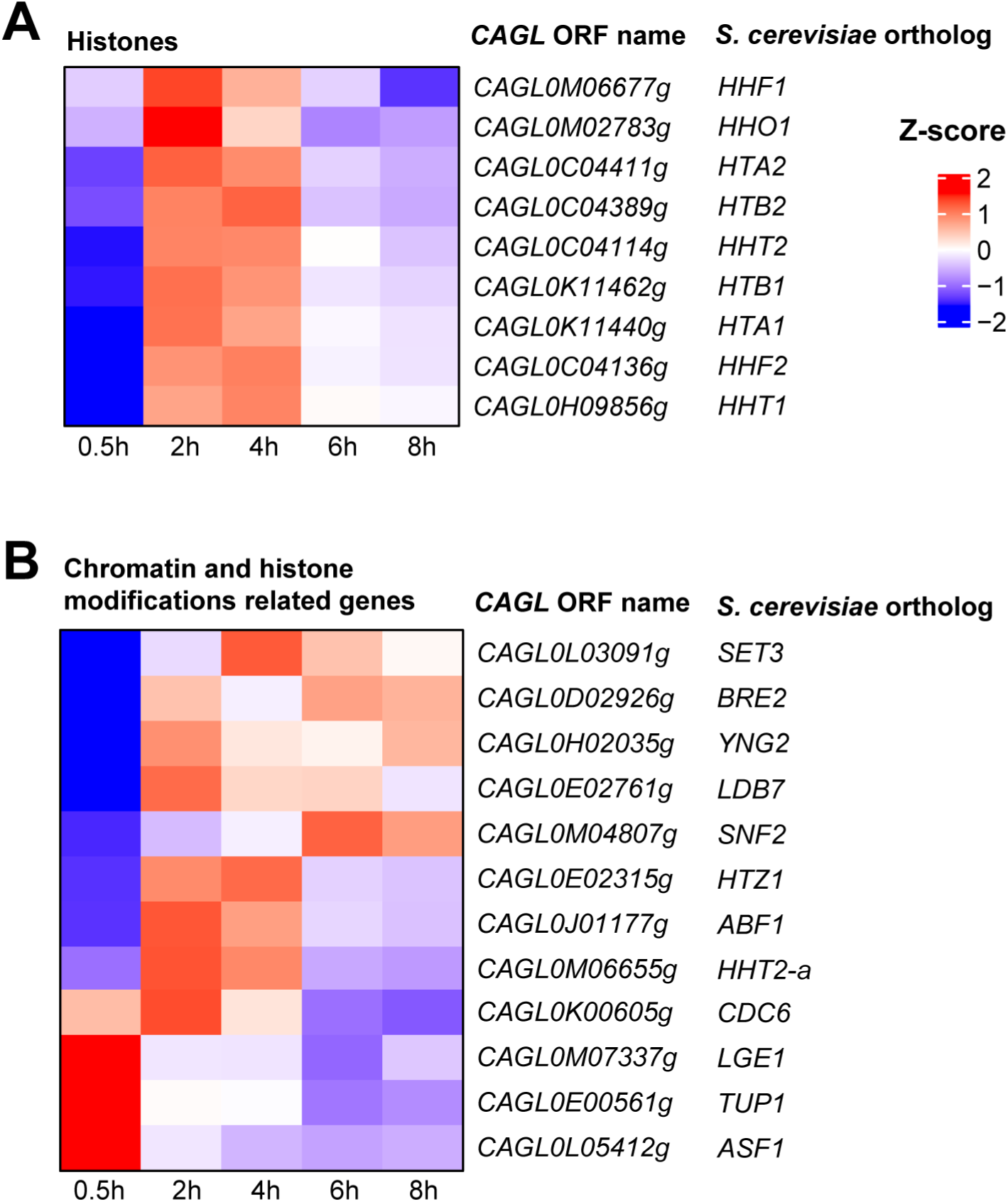
Genes encoding histones or proteins involved in chromatin modification and remodelling are transcriptionally induced in *C. glabrata* during the early stages of macrophage infection. **(A&B)** Heatmaps showing the expression pattern for (A) histone H2A, H2B, H3, and H4 genes and putative (B) chromatin and histone modifiers genes during THP-1 macrophage infection. The colour scale represents the Z-score of the normalized RNAPII ChIP-seq signal.

**Figure 1-figure supplement 5:**
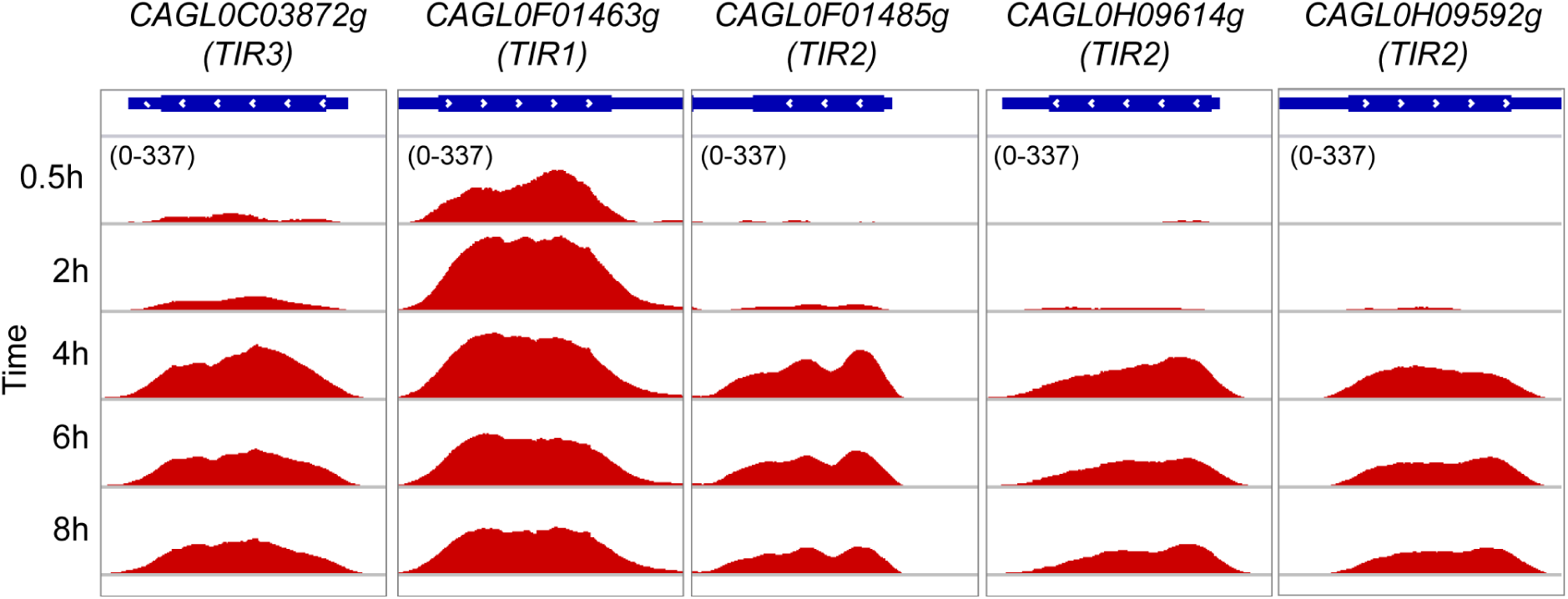
TIR family genes for sterol uptake displaying co-expression during macrophage infection. Genome browser screenshots showing RNAPII ChIP-seq profile on putative TIR family genes for sterol uptake during THP-1 macrophage infection.

**Figure 1-figure supplement 6:**
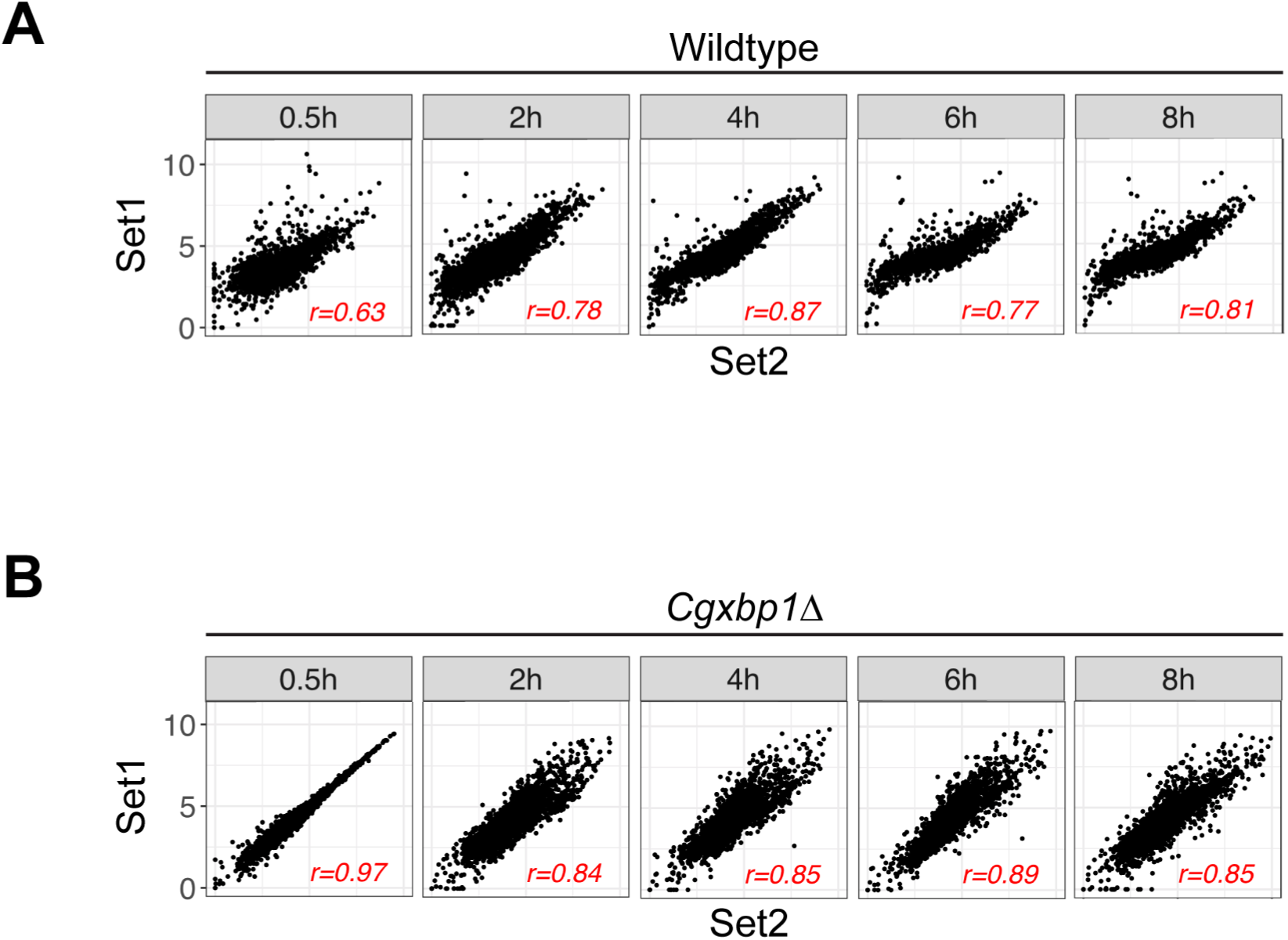
Correlation between independent biological repeats of the RNAPII ChIP-seq experiment for wildtype and the *Cgxbp1*Δ mutant. **(A&B)** Scatterplots showing RNAPII ChIP-seq signals between two independent biological repeats for (A) wildtype and (B) the *Cgxbp1*Δ mutant at the indicated times. The correlation coefficient (r) for each comparison is presented.

**Figure 2-figure supplement 1:**
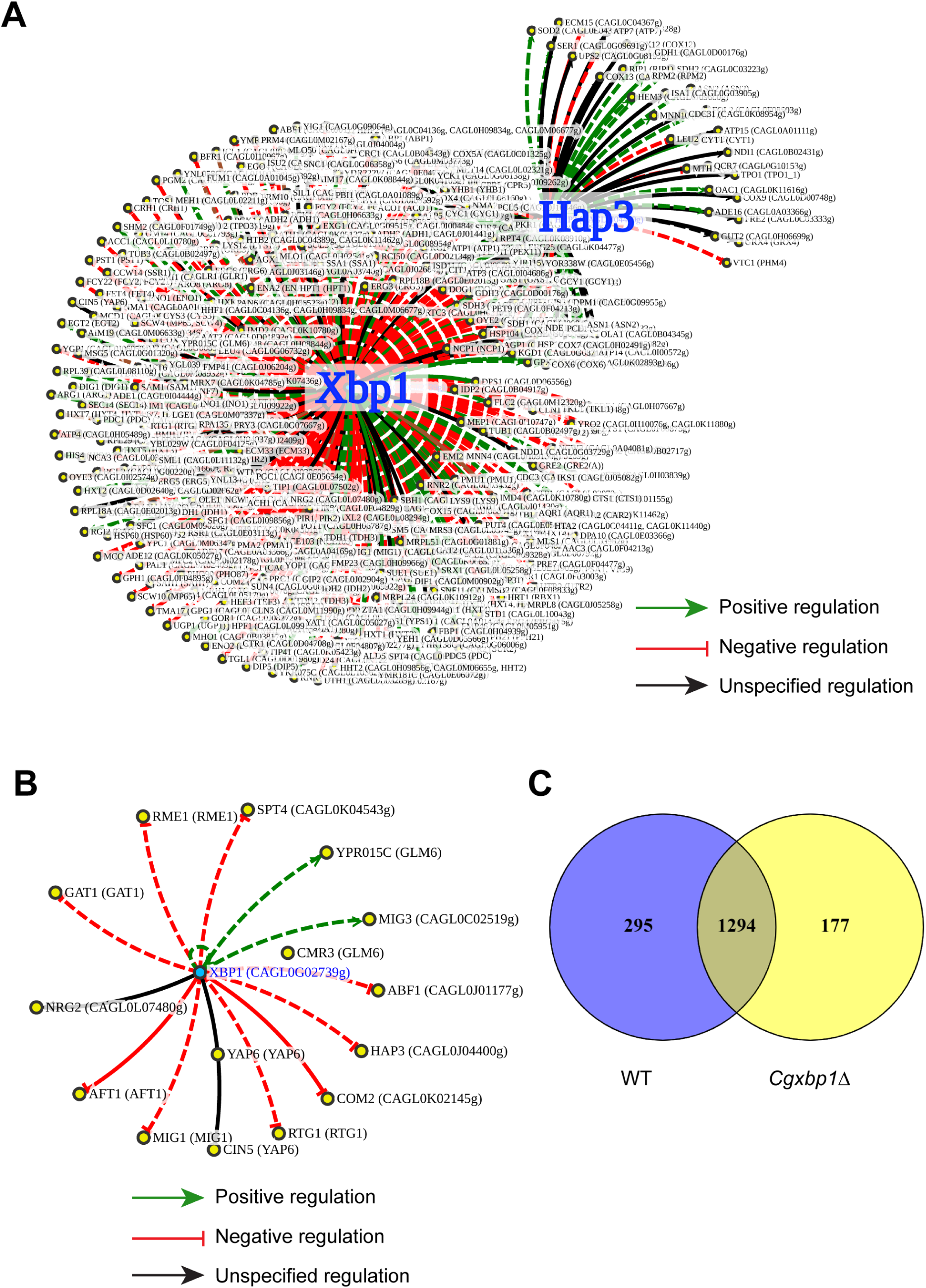
CgXbp1 is a key transcription regulator of the temporal transcriptional response of *C. glabrata* during macrophage infection. **(A)** A regulatory network of Xbp1 and Hap3 for the orthologue of macrophage infection-induced genes in *S. cerevisiae* based on published regulatory information available on the PathoYeastract database. Green, red, and black arrows indicate positive, negative, and unspecified regulation, respectively. Solid and dashed lines represent DNA binding or expression-based evidence, respectively. **(B)** A regulatory network of *CgXBP1* for a subset of infection-induced TF genes based on published regulatory information available on the PathoYeastract database. **(C)** Venn diagram of actively transcribing genes in wildtype and the *Cgxbp1*Δ mutant during macrophage infection.

**Figure 3-figure supplement 1:**
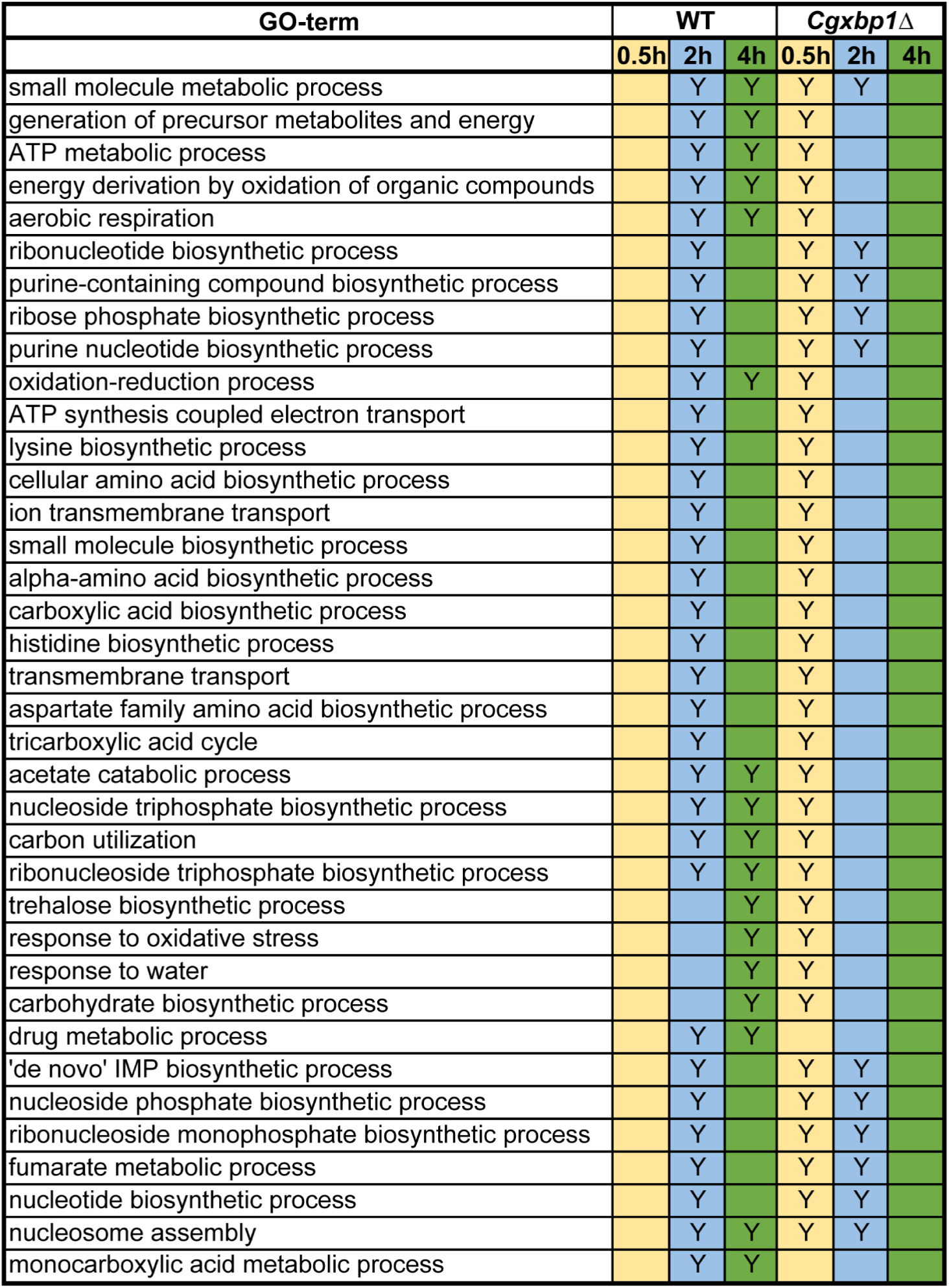
CgXbp1 is essential for the chronological transcriptional response of *C. glabrata* during macrophage infection. A table summarising the enrichment of GO-terms among the transcribed *C. glabrata* genes at the indicated time points during THP-1 macrophage infection by wildtype and the *Cgxbp1*Δ mutant. “Y” indicates a statistically significant enrichment (P value < 0.05) in a given GO term, while a blank box means no significant enrichment.

**Figure 4-figure supplement 1:**
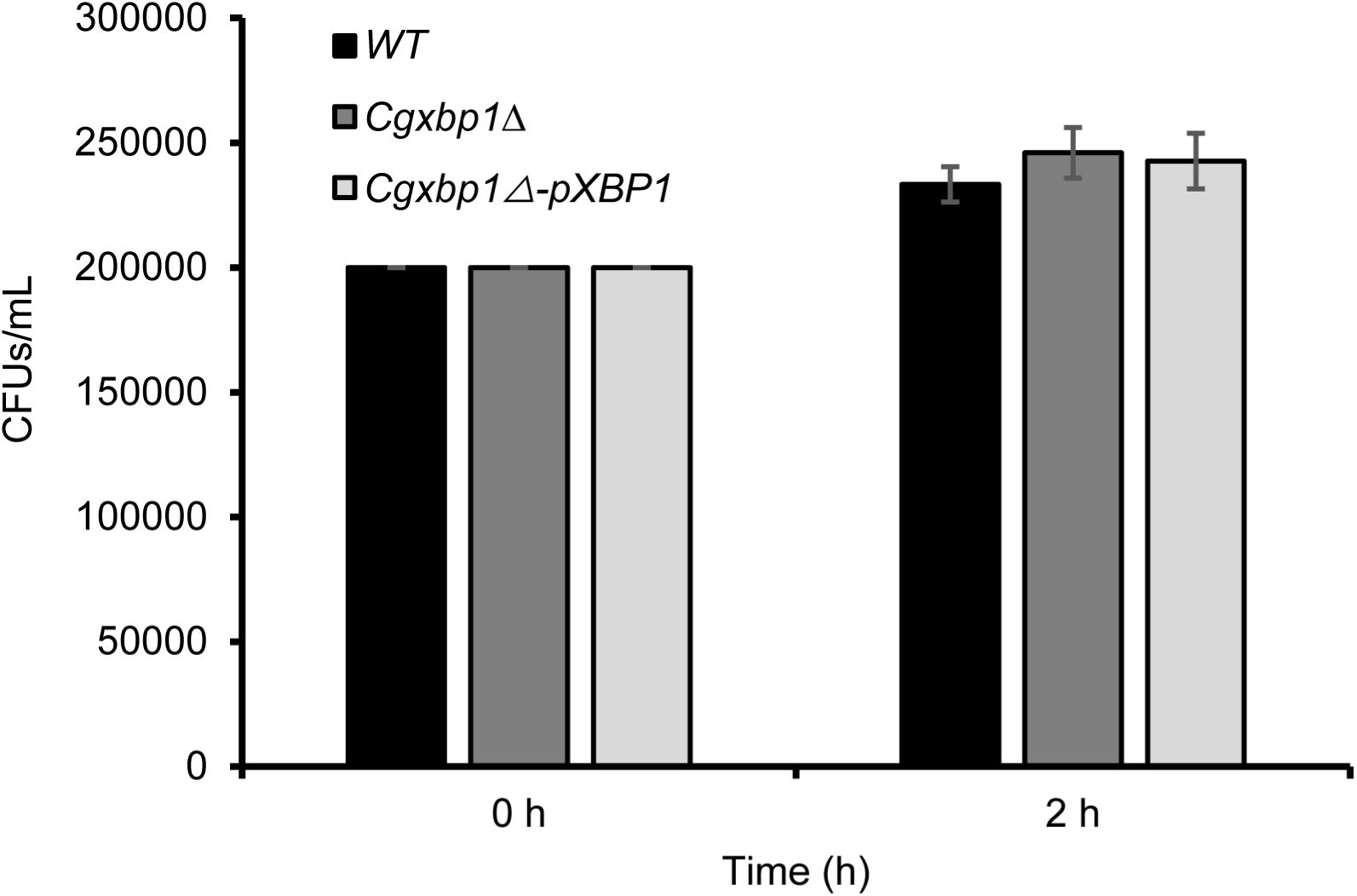
Phagocytosis rate is not affected by CgXbp1 deletion. Bar diagram displaying colony forming units obtained before macrophage infection and 2 h post-infection. Error bars represent mean ± standard deviation from three independent experiments.

**Figure 5-figure supplement 1:**
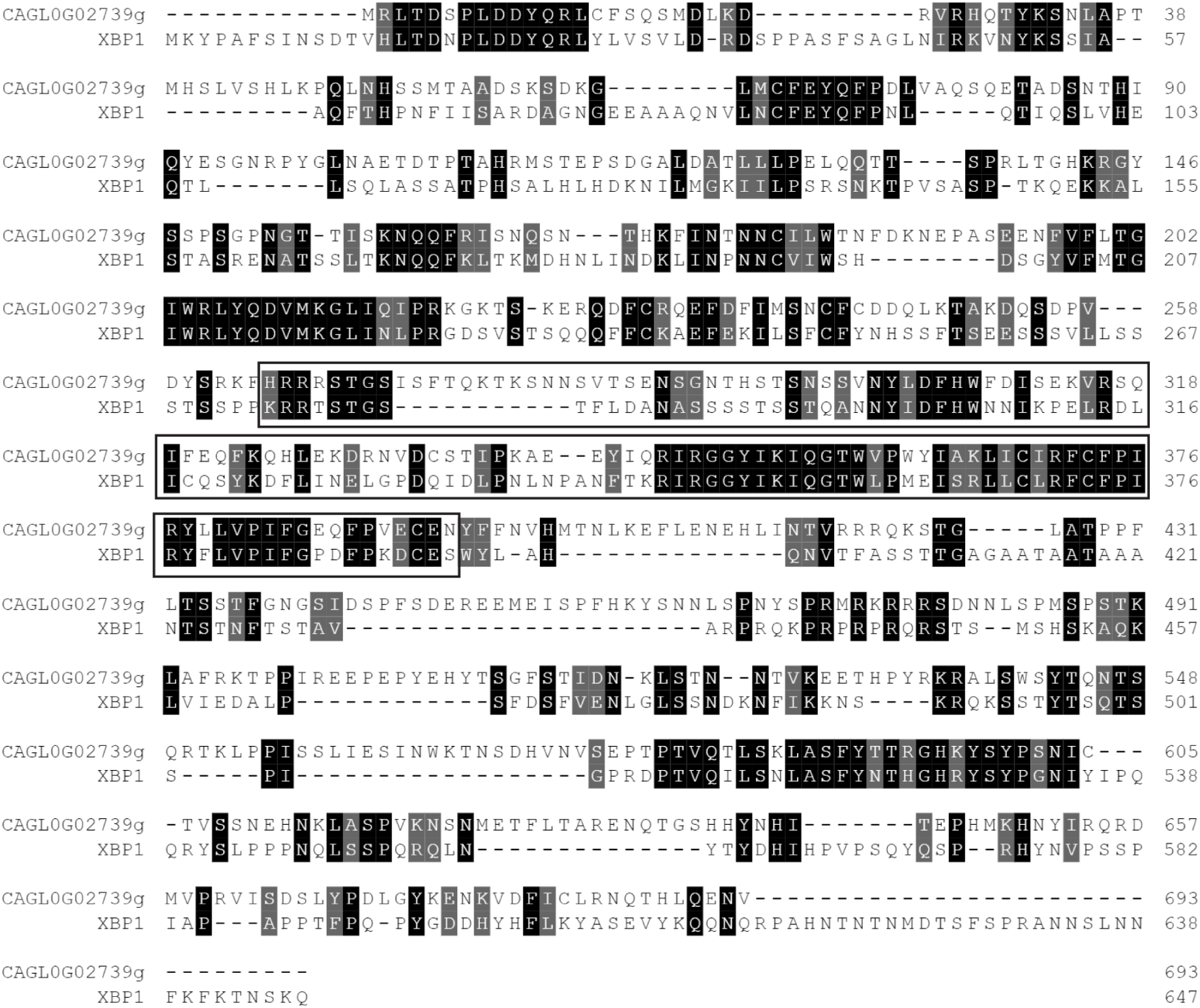
Protein sequence comparison between CgXbp1 and ScXbp1. Pairwise sequence alignment of amino acid sequences of *C. glabrata* ORF *CAGL0G02739g* and *S. cerevisiae* Xbp1. Black and grey shaded amino acid represents identical and similar amino acids, respectively. Boxed amino acid sequence marks the DNA binding domain.

**Figure 5-figure supplement 2:**
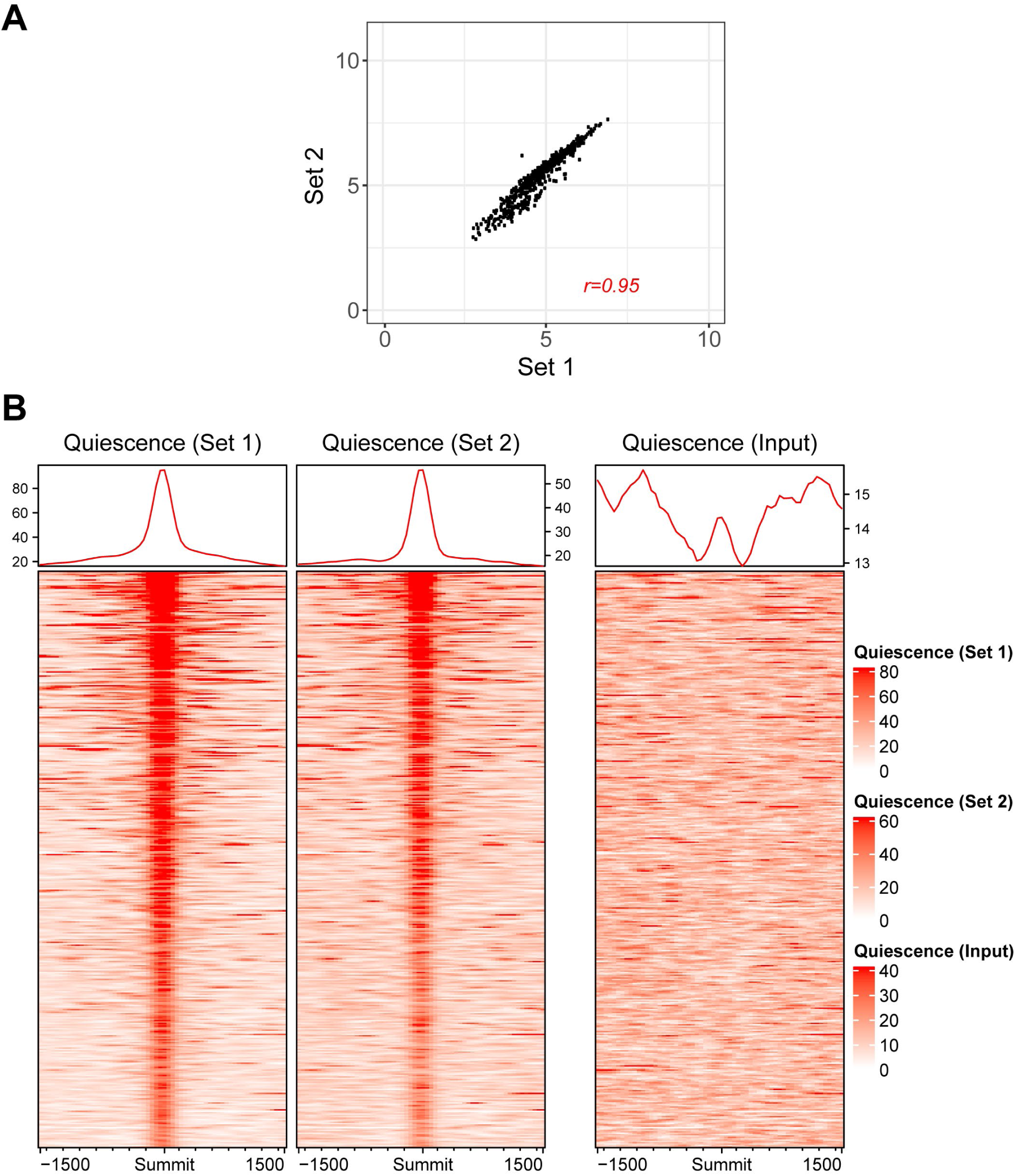
Biological replicates from two independent experiments for CgXbp1^MYC^ ChIP-seq displayed a good correlation. **(A)** Scatterplot showing correlation of ChIP-seq signal intensity at CgXbp1^MYC^ peak summits and 200 bp flanking regions between two independent biological repeats during quiescence phase**. (B)** Heatmap showing ChIP-seq signals on CgXbp1^MYC^ peak summits and 1500 bp flanking regions for two independent biological repeats and the input control.

**Figure 5-figure supplement 3:**
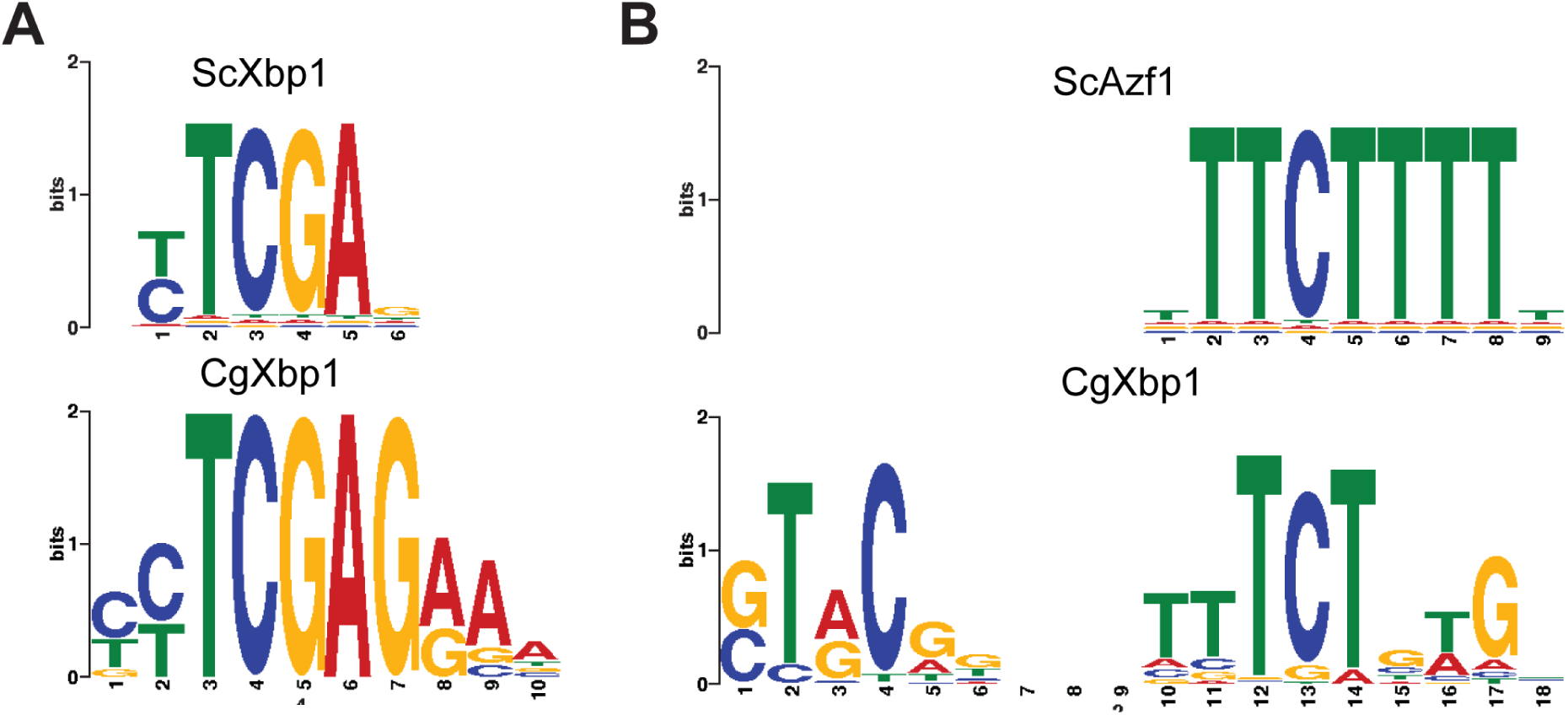
The two enriched motifs identified from CgXbp1 binding sites show good homology with *S. cerevisiae* Xbp1 and Azf1 DNA binding recognition motifs. **(A&B)** TOMTOM suite identifies two motifs in *S. cerevisiae* transcription factors, Xbp1 (A) and Azf1 (B), conserved and similar to CgXbp1.

## Supplementary Files

**Supplementary File 1** - List of actively transcribing genes in wildtype *C. glabrata* upon macrophage infection

**Supplementary File 2** - List of GO-terms enriched from temporally induced genes in wildtype *C. glabrata* in response to macrophage infection

**Supplementary File 3** – Gene regulatory associations between indicated TFs and the macrophage infection-induced genes reported in the PathoYeastract database

**Supplementary File 4** – List of orthologues for the macrophage infection-induced TF and non-TF genes of *C. glabrata* previously shown to have a regulatory association with Xbp1 or Hap3 in *S. cerevisiae* obtained from the PathoYeastract database

**Supplementary File 5** – List of actively transcribing genes in *Cgxbp1*Δ mutant upon macrophage infection

**Supplementary File 6** - List of GO-terms enriched from temporally induced genes in *Cgxbp1*Δ in response to macrophage infection

**Supplementary File 7** - MACS2 output metadata displaying the coordinates of CgXbp1 binding sites during quiescence phase

**Supplementary File 8** - List of CgXbp1 target genes displaying CgXbp1 binding during quiescence phase

**Supplementary File 9** - List of CgXbp1 target genes with two consensus DNA binding motifs

**Supplementary File 10** - List of GO-terms for biological processes enriched from CgXbp1 targets in quiescence phase

**Supplementary File 11 –** List of TF-encoding genes with CgXbp1 binding at their promoters

## Additional files

**Figure 5-Source data** – Western blot original image files used in Figures 5A & B

